# eQTLs identify regulatory networks and drivers of variation in the individual response to sepsis

**DOI:** 10.1101/2023.09.22.558983

**Authors:** Katie L. Burnham, Nikhil Milind, Wanseon Lee, Andrew J. Kwok, Eddie Cano-Gamez, Yuxin Mi, Cyndi G. Geoghegan, GAinS Investigators, Stuart McKechnie, Nicole Soranzo, Charles J. Hinds, Julian C. Knight, Emma E. Davenport

**Affiliations:** Wellcome Sanger Institute, Wellcome Genome Campus, Hinxton, UK; University of Cambridge, Cambridge, UK; Wellcome Centre for Human Genetics, University of Oxford, Oxford, UK; Oxford University Hospitals NHS Foundation Trust, Oxford; Centre for Translational Medicine & Therapeutics, William Harvey Research Institute, Faculty of Medicine & Dentistry, Queen Mary University of London, London, UK; Chinese Academy of Medical Science Oxford Institute, University of Oxford, Oxford, UK

## Abstract

Sepsis is a clinical syndrome of life-threatening organ dysfunction caused by a dysregulated response to infection, for which disease heterogeneity is a major obstacle to developing targeted treatments. We have previously identified gene expression-based patient subgroups (Sepsis Response Signatures: SRS) informative for outcome and underlying pathophysiology. Here we aimed to investigate the role of genetic variation in determining the host transcriptomic response and to delineate regulatory networks underlying SRS. Using genotyping and RNA-seq data on 638 adult sepsis patients, we report 16,049 independent expression (eQTLs) and 32 co-expression module (modQTLs) quantitative trait loci in this disease context. We identified significant interactions between SRS and genotype for 1,578 SNP-gene pairs, and combined transcription factor (TF) binding site information (SNP2TFBS) and predicted regulon activity (DoRothEA) to identify candidate upstream regulators. These included HIF1A and CEBPB, which were associated with progenitor and immature neutrophil subsets respectively, further implicating glycolysis and emergency granulopoiesis in SRS1. Overall, these approaches identified putative mechanistic links between host genetic variation, cell subtypes, and the individual transcriptomic response to infection. Understanding the regulatory networks underlying patient heterogeneity provides additional information for developing immunomodulatory treatments and a personalised medicine approach to treating sepsis.

## Introduction

Sepsis is a clinical syndrome defined as organ dysfunction resulting from a dysregulated immune response to infection^1^, which causes an estimated 11 million deaths per year worldwide^2^. The sepsis response involves concurrent proinflammatory and immunosuppressive mechanisms, the extent and impact of which varies considerably both between and within patients over time^3,4^. The complexity and heterogeneity of the host sepsis response have limited attempts to develop targeted treatments^5^, which require better understanding of individual responses to infection and the predominant underlying mechanisms that drive organ dysfunction^6^. We, and others, have stratified sepsis patients by clinical and molecular measures to define more homogeneous subgroups^7–11^. These subgroups are hypothesised to have some distinct underlying pathophysiological mechanisms, and thus could be leveraged to identify the key drivers acting in these different contexts. For example, we have previously shown that Sepsis Response Signature (SRS) subgroups resolve the majority of transcriptomic variation in sepsis^7^ even accounting for different infection sources^12–14^, and can also be assessed based on a quantitative score (SRSq)^13^. SRS1, and increasing SRSq, identify a group of patients with features of immunosuppression and a higher risk of mortality, but the regulatory determinants and predisposing factors are unclear.

Host genetic background is a plausible driver of variation in the response to infection^15–18^, and in this regard use of genome-wide association studies and rare variant analysis has proved highly informative in severe COVID-19^19–23^. While these approaches have also been applied in sepsis^16,17,24–26^, they have been less successful, again likely due to disease heterogeneity. We hypothesise that the host response to sepsis, as represented by the transcriptomic SRS subgroups, has a polygenic basis with additional environmental modulators, and that elucidation of these underlying regulatory networks will be key in understanding inter-individual variation in disease responses and identifying treatable traits^27^. Genetic factors, notably non-coding variants, may modulate gene expression as an expression quantitative trait locus (eQTL). Importantly, such associations can vary across cell types and states^28–30^, meaning that it is vital to map eQTLs in the context of disease to understand the contribution of regulatory variants to pathogenesis^31^. Additionally, specific instances of transcriptomic regulation may be affected by external factors, resulting in a genotype-by-environment interaction. Identification of a set of eQTLs that are modulated by the same environmental variable can therefore reveal important upstream regulators and pathways through which the perturbation impacts biological function^28,29^.

Here, we apply this approach to sepsis, particularly investigating the influence of genetic variation on transcriptomic subgroups that may represent divergent regulatory networks. The UK Genomic Advances in Sepsis (GAinS) study has recruited a cohort of >1400 patients with sepsis due to community acquired pneumonia (CAP) and faecal peritonitis (FP), admitted to adult intensive care units (ICUs) across the UK. We have generated array genotyping data on 1,168 patients and genome-wide gene expression data on 1,043 unique patients^7,12,13^. This provides an opportunity to identify genetic drivers of gene expression in a disease context with high power to detect interaction effects related to patient phenotypes. We find evidence of genetic variation associated with widespread transcriptomic differences in the sepsis response and leverage this to identify putative key regulators of SRS.

## Results

### Evidence for contribution of genetic variation to SRS

We have assigned SRS group membership to 997 GAinS sepsis patients with both genotyping and blood gene expression data (from qPCR, microarray or RNA-sequencing^7,12,13^) collected at multiple time points during the first five days of ICU admission (Figure 1a). We first aimed to estimate the overall contribution of common genetic variation to the SRS phenotype using paired genotyping and, as SRS status for a given patient may change over time with disease natural history, whether a patient had ever been assigned to SRS1 during the period of observation. We estimated the heritability^32^ as 57% (±28%, p=0.019, n=440 “SRS1ever” vs 557 “SRS1never” patients), supporting the hypothesis that common variants contribute substantially to SRS during critical illness (Supplementary Figure 1).

**Figure 1:**
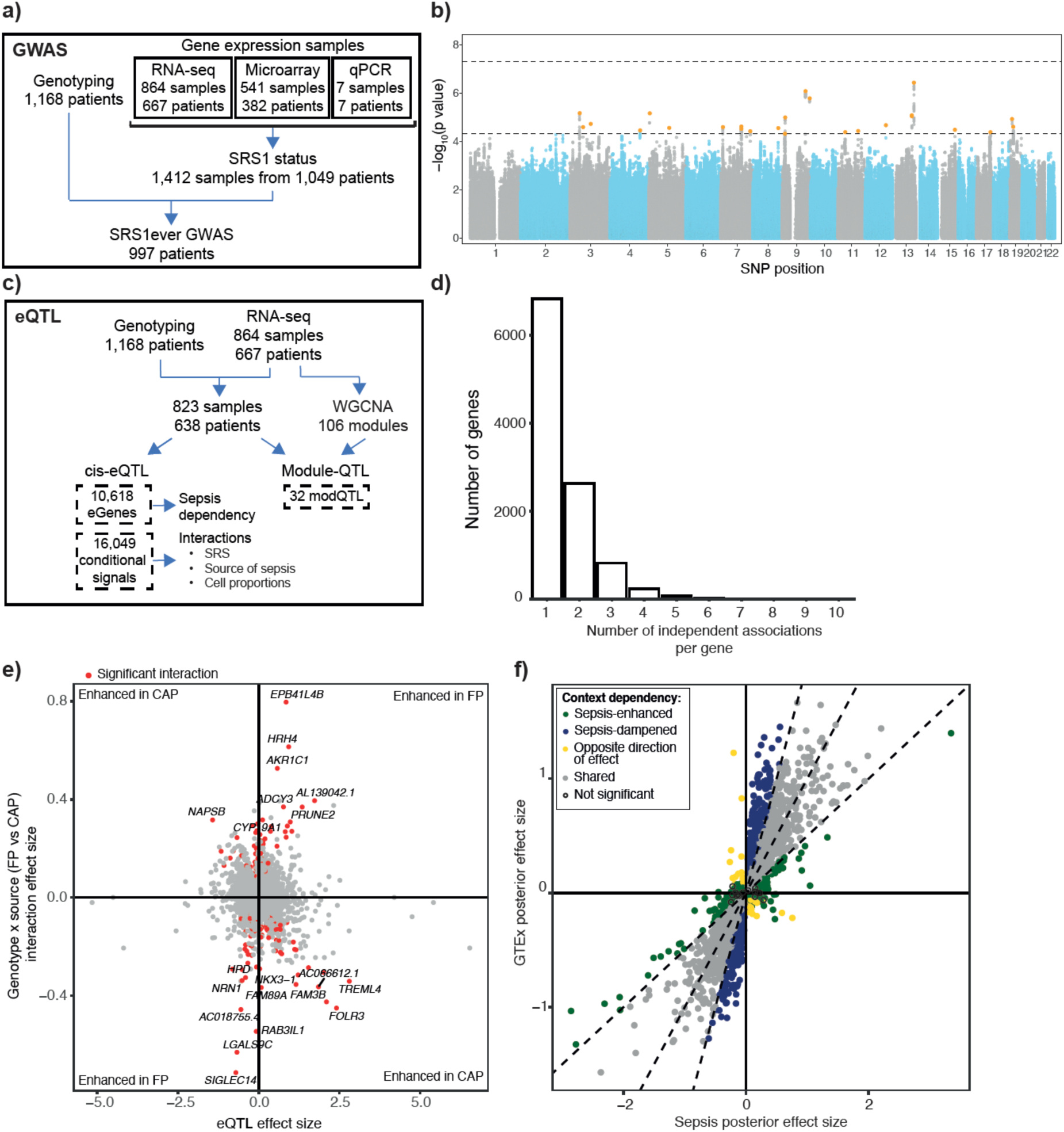
Genetic variants associated with gene expression in the context of sepsis. **a)** Schematic of cohort design for SRS1ever genome-wide association study (GWAS) using all patients with genotyping data and at least one gene expression time point for SRS assignment. **b)** Common SNPs (MAF≥1%) were tested for association with the SRS1ever vs never phenotype. Manhattan plot showing −log10(p-value) for each variant plotted against its genomic position. The most significant SNP in each locus is highlighted in orange. **c)** Schematic of cohort design for eQTL (*cis*-eQTL and coexpression module QTL) analysis using all samples from patients with genotyping data and RNA-seq data. Coexpression modules were defined using the full RNA-seq dataset. **d)** Histogram showing the distribution of the number of independent signals detected through conditional analysis for each eGene. **e)** eQTL interactions with source of sepsis (CAP or FP). Each point represents an independent eSNP-eGene pair, with the interaction effect size plotted against the genotype effect. eQTLs with bigger effects in FP compared to CAP are therefore found in the top right and bottom left quadrants. Red colour indicates a significant interaction between genotype and source of sepsis (FDR<0.05), with the most significant results labelled with the eGene name. **f)** Sepsis-dependent eQTL effects identified with mashr. Each point represents a lead SNP-eGene pair from the first pass sepsis eQTL mapping that was also tested for whole blood eQTL in the European subset of GTEx. Posterior effect sizes estimated by mashr are plotted for GTEx against sepsis, and eQTLs are categorised as shared or sepsis-dependent based on the difference between these estimates (significance, effect sizes differing by a factor of 0.5).

We proceeded to carry out a genome-wide association study (GWAS) for SRS1ever vs never. This identified 155 variants in 25 loci reaching the genome-wide suggestive p-value threshold of 5×10^-5^, henceforth referred to as “SRS1 GWAS SNPs” (Figure 1b, Supplementary Figure 2, Supplementary Table 1). These include SNPs previously reported to be associated with monocyte, lymphocyte, and platelet counts, clotting factors, and lung function^33,34^ (Supplementary Table 2).

### Expression quantitative trait loci in sepsis patients

To identify genetic regulators of gene expression, we then mapped *cis*-eQTLs using a random intercept linear mixed model^28^ (823 samples from 638 sepsis patients with bulk RNA-seq and genotyping, Figure 1c, Supplementary Figure 3, Methods) and identified significant associations for 10,618 eGenes (Supplementary Table 3). Given the importance of the MHC region in immunity^35^, we ensured accurate quantification of HLA gene expression and eQTL identification through re-mapping to personalised references (Methods). Through conditional analysis we found 16,049 independent associations in total (Supplementary Table 4), with multiple signals for 3,788 eGenes (35.7% of all eGenes, median of 1 and a maximum of 10 signals for each gene) (Figure 1d). As previously reported in healthy cohort studies^36^, we observed that the eSNPs underlying primary signals were more common and located closer to the transcription start site of the associated gene than were secondary/tertiary+ signal variants (Supplementary Figure 4). Five sepsis eQTLs involved SRS1 GWAS SNPs (Supplementary Table 2). Of these, two eQTL signals in the same region (*OCEL1,* a predicted membrane component^37^, and *NR2F6*, an immune checkpoint which suppresses adaptive responses^38^) showed evidence of colocalisation^39^ with the same GWAS locus (PP4 90.3% and 89.9% respectively) indicating the same causal variant is driving these signals (Supplementary Figure 5).

We have previously described eQTLs in a smaller microarray dataset of patients with sepsis due to CAP (n=240, of which 134 individuals overlap with the RNA-seq cohort, Methods)^7^. The previously reported eQTL effect sizes for SNP-gene pairs assayed in both datasets are significantly correlated with this new dataset (Pearson’s r=0.72, Supplementary Figure 6, Supplementary Table 5), but we also identify more than twice the number of eGenes, likely due to the larger sample size and the greater sensitivity of RNA-seq for detecting expression. This expanded cohort also encompasses a broader range of patients, including abdominal sepsis, meaning some of the additional eQTL detected here may be context-dependent. We tested each of the independently associated lead SNPs for interaction effects with the source of sepsis (CAP or FP, 12,663 SNP-gene pairs with ≥2 minor allele homozygotes in each group tested, Methods). We identified 166 significant interaction effects (FDR<0.05), more than expected by chance (permutation p-value <0.01, Supplementary Figure 7, Supplementary Table 6), of which roughly half (n=88) had stronger effects in CAP (Figure 1e). The eGenes involved were enriched for the Reactome term “Biological oxidations” (FDR=0.0033) and were members of a subnetwork connected by hub genes *APP*, A*KT1 and ABCC1* (Methods).

To explore sepsis-dependency of observed eQTLs, we used GTEx and eQTLgen, two well-characterised blood eQTL datasets from healthy donors^40,41^. As expected, the majority of effects were highly correlated between health and sepsis (Pearson’s r=0.898, 0.803) but we did find *cis*-eQTLs in sepsis not detected in either healthy dataset (38.7% and 17% of our lead SNP-gene pairs were called non-significant in GTEx and eQTLgen respectively) (Supplementary Figure 8). Given this evidence, we further quantified this context dependency by comparing sepsis eQTL effect sizes to GTEx, a similarly sized bulk RNA-seq cohort, with mashr^42^ (Methods). We categorised our eQTLs significant in sepsis as “shared” (n=5,943) or “context-dependent” (n=2,179, Figure 1f). Specifically, we identified 854 signals with bigger effects and/or only significant in sepsis (“sepsis-enhanced”), 1,272 with bigger effects in healthy volunteers (“sepsis-dampened”) and 53 eQTL that were significant in both GAinS and GTex but had opposite directions of effect (Supplementary Table 7). We found no significantly enriched Reactome pathways within these eGene subsets. However, sepsis-enhanced eQTLs differ significantly from those shared with GTEx, with the eSNPs involved having a lower MAF (median 11.5% vs 12%, Mann-Whitney p=0.00015), and further away from the TSS of their target gene (median |distance|=52.1kb vs 27.6kb, Mann-Whitney p=2.5e-27, Supplementary Figure 9).

### eQTL effects vary between sepsis patients

We proceeded to investigate the distinct regulatory networks underlying the SRS groups by identifying differential genetic regulation of gene expression between samples assigned to SRS1 and non-SRS1. We found a significant interaction between genotype and SRS status in 1,578/12,959 of SNP-gene pairs (12%), with ⅔ of these having a more pronounced effect on gene expression in SRS1 (1,064 magnified in SRS1 vs 514 dampened; Figure 2a, Supplementary Figure 10; Supplementary Table 8). eGenes with a magnifying SRS interaction were enriched for the Reactome term “synthesis of PA [phosphatidic acid]” (FDR=0.0069). All SRS interaction eGenes were enriched for genes differentially expressed between SRS groups^13^ (FET p-value 4.59 x 10^-4^), with 628 magnifiers and 105 dampeners involving genes that were significantly upregulated in SRS1 (enrichment of magnifiers in upregulated genes FET p-value=5.3×10^-63^). We noted that sepsis-enhanced eGenes were significantly enriched for SRS interactions (FET p-value=0.03), indicating that genetic regulation of some genes may be modulated on a spectrum from health to SRS2 to SRS1. This is illustrated by a subset of sepsis-enhanced eGenes where the effect of genotype also increases continuously with SRSq score, a quantitative measure for SRS^13^; for example, *FAM89A*, a gene previously highlighted as a potential biomarker for paediatric bacterial infection^43^ (Figure 2b).

**Figure 2:**
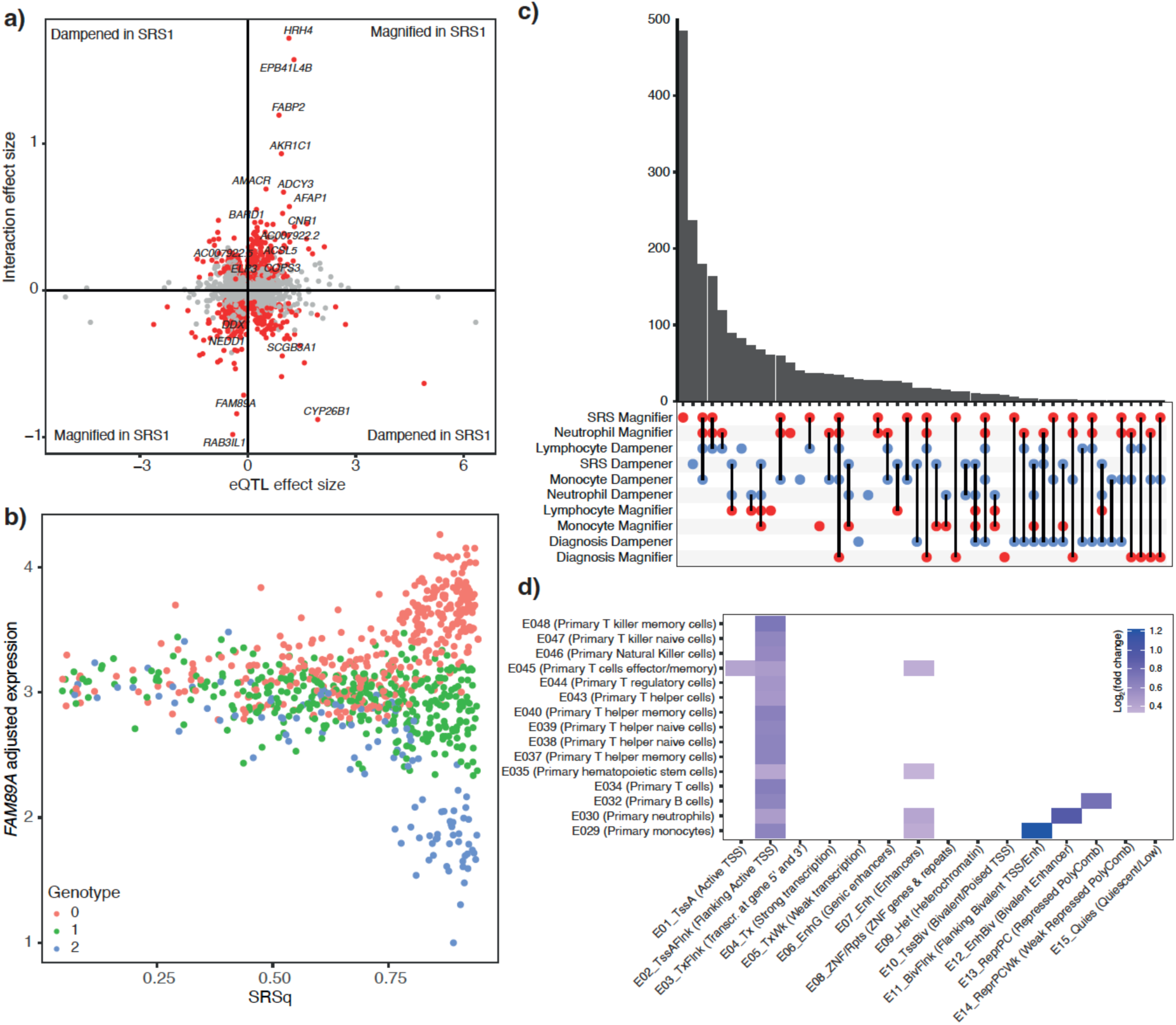
Genotype-by-environment interactions find widespread variation in eQTL effects across sepsis patients. **a)** eQTL interactions with SRS1 status. Each point represents an independent eSNP-eGene pair, with SRS interaction effect size plotted against the genotype effect. eQTLs with bigger effects in SRS1 compared to non-SRS1 are therefore found in the top right and bottom left quadrants. Red colour indicates a significant interaction between genotype and SRS1 status (FDR<0.05), with the most significant results labelled with the eGene name. **b)** An exemplar sepsis-enhanced eQTL that also has a significant positive interaction with SRS1 status. Gene expression residuals plotted against SRSq, with point colour indicating number of copies of the minor allele. **c)** UpSet plot showing sharing of magnifying (red) and dampening (blue) eQTL interaction effects between environmental variables tested. **d)** Enrichment of eSNPs for eQTLs with an SRS interaction in different chromatin states from the Roadmap Epigenomics Core 15-state Genome Segmentation annotations^46^ across relevant cell types, compared to eSNPs for eQTLs without a significant SRS interaction (only significant enrichments shown [FDR<0.05]).

As previously demonstrated in healthy cohorts^44,45^, we identify many eQTLs with putative cell type-specific effects by adding genotype-cell proportion interaction terms to our eQTL model. Specifically, we found 1,073, 1,013, and 608 eQTLs with significant interactions (FDR<0.05) with measured neutrophil, lymphocyte, and monocyte proportions respectively (Supplementary Figure 11, Supplementary Table 9). The larger number of interactions detected in neutrophils and lymphocytes may be due to those cell types being more prevalent and having greater variance across sepsis patients. As cell proportions are necessarily related, we observe highly correlated and reciprocal effects across interaction types; for example, eQTLs whose effect is amplified in the context of high neutrophil proportions (“neutrophil magnifiers”) generally also had a larger effect size with lower lymphocyte proportions (“lymphocyte dampeners”) (Figure 2c).

We observed that SRS interaction eSNPs were enriched (one-tailed binomial test FDR <0.05) in neutrophil and monocyte enhancers and flanking TSS regions across primary immune cells^46^ (Figure 2d, Methods), indicating the particular contribution of myeloid cells to the transcriptomic differences between SRS groups. Given recent findings that disrupted granulopoiesis is a key feature of the SRS1 subgroup^14^, we were interested to note that SRS1 magnifiers showed a large overlap with neutrophil magnifiers (496/1064) (Figure 2c). One such SRS and neutrophil magnifier eQTL involved an SRS GWAS SNP (rs891204), associated with reduced expression of *OCEL1*.

### Upstream regulatory drivers of SRS

We then used the SRS interaction eQTLs to identify putative upstream drivers of the differential host response to sepsis. We hypothesised that variation in the impact of regulatory variants could be mediated by changes in the activity of a transcription factor whose binding is affected by the eSNPs. We therefore first identified instances where each of 127 human transcription factor binding motifs were interrupted or introduced by sepsis eSNPs (or their LD proxies) using SNP2TFBS^47,48^ (Figure 3a). We found this was more common when the eQTL had an SRS interaction (median 5 vs 4 binding sites introduced/interrupted per eGene, Wilcoxon p-value=2.68×10^-7^). For each TF motif, we then classified eGenes as having at least one or no binding sites altered. We found 59 TF motifs were significantly enriched for alteration among SRS interaction eGenes (Figure 3b, Supplementary Table 10), with the HIF1A-ARNT motif having greatest enrichment. This was significantly more motifs than expected by chance (p<0.001), as computed by permuting eGene interaction status 1000 times and repeating the enrichment analysis (Supplementary Figure 12).

**Figure 3:**
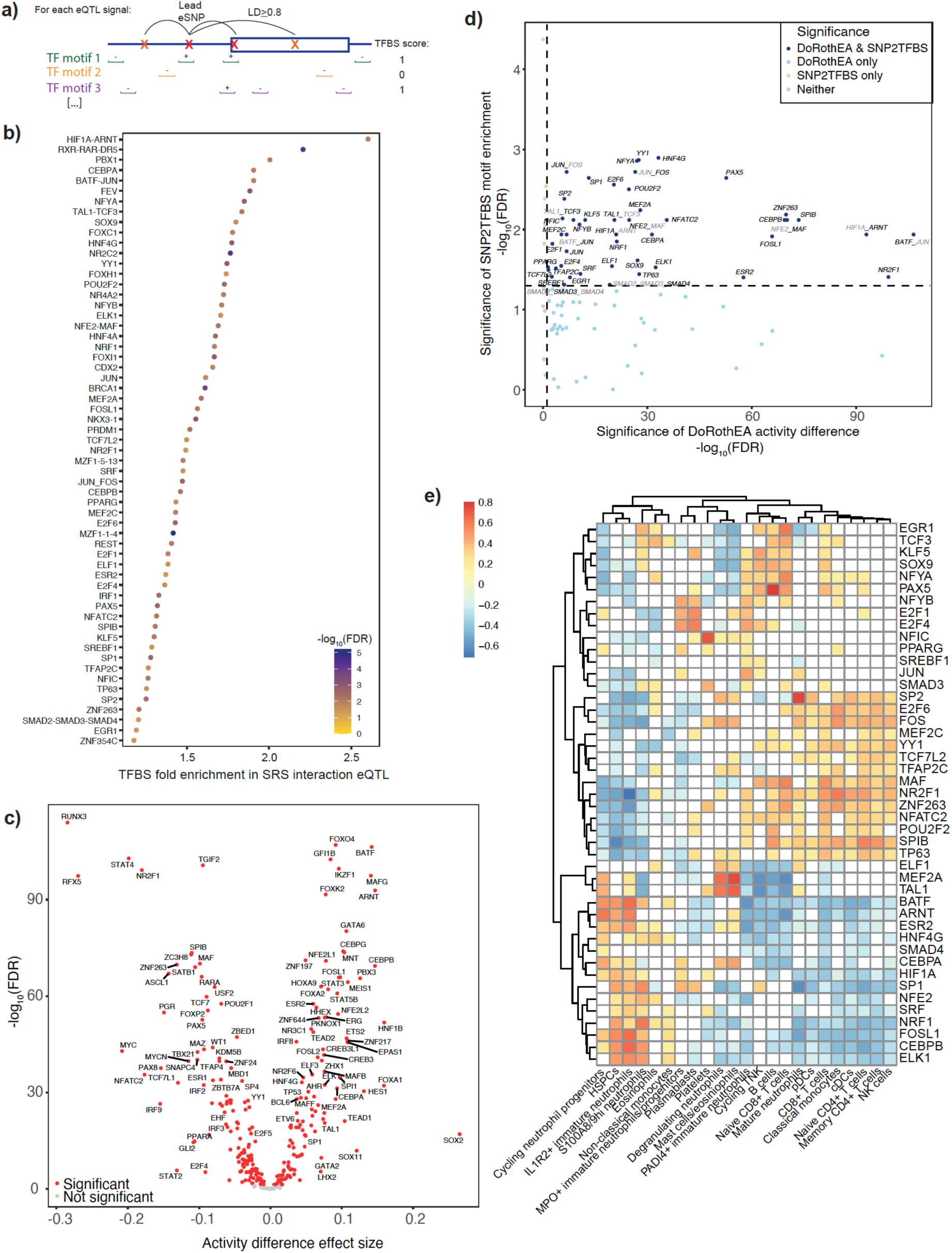
Identification of putative driver transcription factors for SRS from eQTL interactions. **a)** Schematic of strategy to identify transcription factor binding sites (TFBS) enriched in SRS interaction eQTLs. For each of 12,959 eQTL signals tested for an SRS interaction effect, query SNPs were defined as those in LD (r^2^≥0.8) with the lead SNP. We then identified instances where binding motifs for 127 human TFs were interrupted or introduced by these query SNPs using SNP2TFBS. Each independent eQTL signal was scored for each motif as having ≥1 or 0 binding sites altered by the signal SNP or its LD proxies. We then tested for enrichment of each TF motif amongst eQTLs with a significant interaction effect compared to eQTLs with no significant interaction using a one-sided Fisher’s exact test. **b)** Transcription factor binding sites identified with SNP2TFBS enriched in eQTLs with an SRS interaction vs no interaction, with point colour indicating significance. **c)** Volcano plot showing comparison of inferred transcription factor activity between SRS1 and non-SRS1 samples. Each point represents one TF, with adjusted p-value plotted against effect size estimated from a linear mixed model. Red colour indicates significance. **d)** Adjusted p-values from b) are plotted against those from c) to highlight TFs with both enriched binding sites in SRS interaction eQTLs and differential activity between SRS. **e)** Heatmap showing Spearman correlation between estimated cell proportions and inferred TF activity for the first sample available per patient. White colour indicates a non-significant correlation (FDR>0.05), red a positive correlation and blue a negative correlation.

However, these predicted binding sites may not be occupied *in vivo*, and experimental binding data is currently limited to small numbers of factors and contexts. We therefore calculated transcription factor activity scores in each sample to pinpoint regulators predicted to vary by SRS (Methods)^49,50^. We found that 253/288 factors had significantly differential inferred activity between SRS groups (Figure 3c, Supplementary Table S11). Given the previously reported extensive differential gene expression between SRS groups^7,13^, this is not surprising, but we confirmed that it was more than expected by chance by permuting SRS status and recalculating differential activity (p<0.001, Supplementary Figure 13).

Of the TFs with differential activity, 45 were also enriched in the SNP2TFBS analysis (41 motifs), indicating that they could be driving the differences in the regulatory landscape between SRS groups (Figure 3d). These included BATF-JUN, ZNF263, HIF1A-ARNT, CEBPB, SPIB, NFE2-MAF and FOSL1. We also utilised cell proportions estimated in each sample using a sepsis single cell reference dataset^14^ to identify relationships between these TFs and 23 specific cell types. The inferred activity of many of the putative driver TFs were weakly correlated with a number of the 23 estimated cell subsets, but some had strong and specific associations, potentially indicating cell type specificity. These included PAX5 with B cells (rho=0.80), SP2 with mature neutrophils (rho=0.76), MEF2A and TAL1 with mast cells/eosinophils (rho=0.75, 0.69) and degranulating neutrophils (0.64, 0.61), NFIC with platelets (0.72), CEBPB, FOSL1 and BATF with IL1R2 immature neutrophils (0.66, 0.65, 0.64)), and ARNT with cycling neutrophil progenitors (0.62) (Figure 3e, Supplementary Table 12).

### Co-expression module QTL mapping

Finally, we aimed to investigate the regulation of gene expression in *trans*. Given our cohort size, we focused on one possible mechanism where a single SNP is associated with the expression of multiple related distal genes via a shared upstream regulator^41^. To increase our power to detect such trans-regulatory networks, we leveraged metrics summarising the expression patterns of co-expressed gene sets, rather than testing each gene individually.

Using the WGCNA package^51,52^ we identified 106 co-expression modules, each comprising 11-1,785 genes (Supplementary Table 13, Supplementary Figure 14), and summarised the expression of each module with its primary eigengene (the first principal component). We found that individual modules were associated with disease phenotypes including SRS, survival and measured cell proportions (Supplementary Figure 15, Supplementary Table 14). Furthermore, these modules are enriched for biological pathways and marker genes for more granular blood cell populations^14,53^ suggesting they capture gene sets associated with specific biological processes (Figure 4a, Supplementary Figure 16, Supplementary Table 15, 16, 17). Finally, to investigate whether these co-expression modules represent sets of co-regulated genes, we tested each module for enrichment of known TF targets from DoRothEA^49^ and found at least one significant TF for 43/106 modules (Supplementary Table 18).

**Figure 4:**
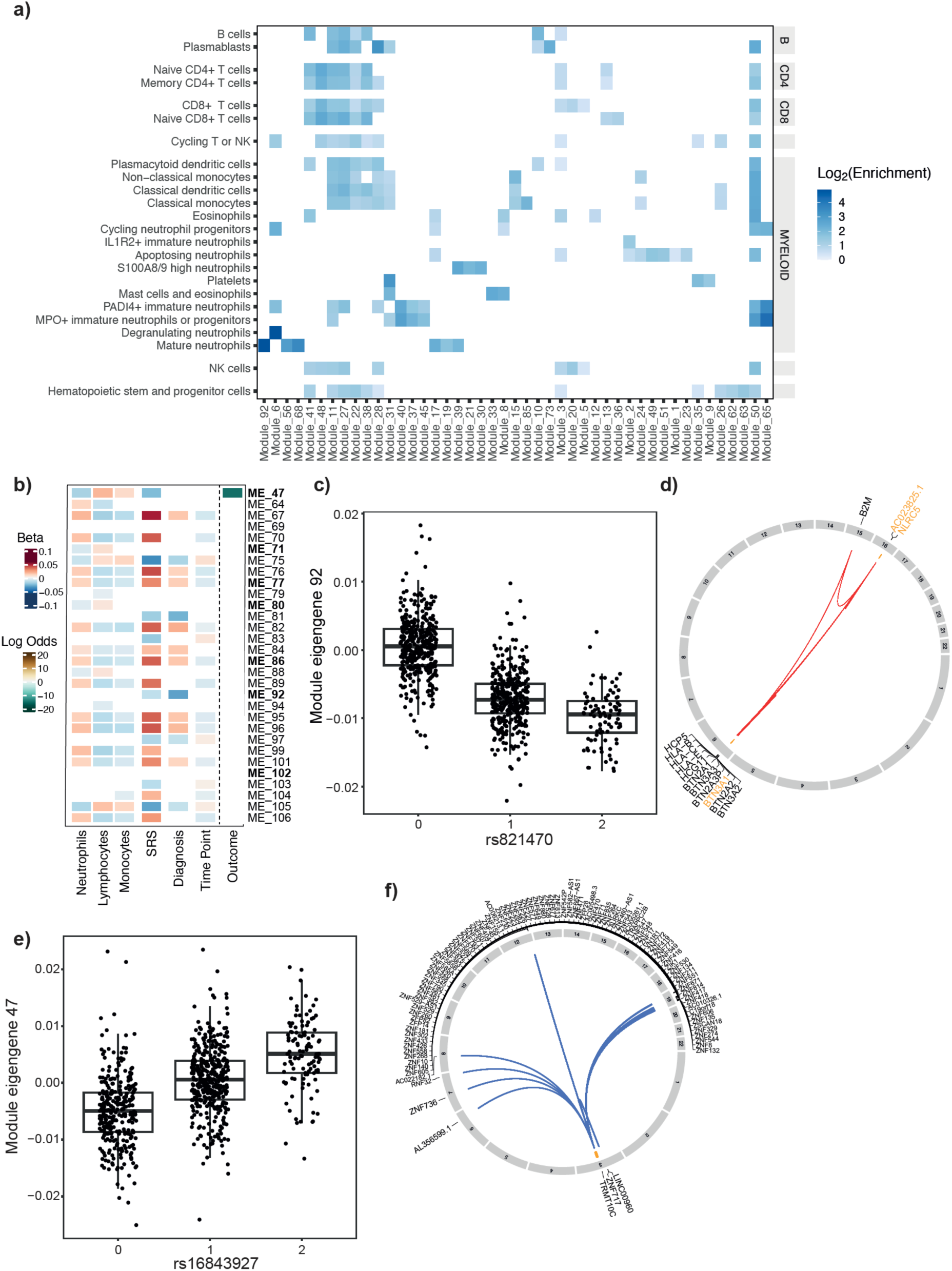
Co-expression modules pinpoint several trans-regulatory networks relevant to sepsis outcomes. **a)** Heatmap showing enrichment of sepsis cell markers in module member genes. Modules were tested for enrichment of leukocyte marker genes identified in a sepsis cohort^14^. Modules shown had significant enrichment for at least one signature. **b)** Heatmap showing significant associations between module eigengenes (MEs) with a modQTL and clinical phenotypes. ModQTLs passing stringent sensitivity analysis are marked with bold font. MEs were tested for differential expression with measured cell proportions, SRS1 status, diagnosis (CAP or FP) and time point (day 1, 3, 5) using a linear mixed model. Association of each ME with survival up to 28 days was tested using a Cox proportional hazards model. **c)** Module 92 eigengene (ME_92) plotted by rs821470 genotype, with partial residuals calculated from the linear mixed model fit. **d)** Circos plot showing the chromosomal locations of the genes contained in module 92 and the lead sepsis eSNPs associated with the ME. Member genes that are *cis*-eGenes for these eSNPs are highlighted in orange. **e)** Module 47 eigengene (ME_47) plotted by rs16843927 genotype, with partial residuals calculated using the linear mixed model fit. **f)** Circos plot showing the chromosomal locations of the genes contained in module 47 and the lead sepsis eSNPs associated with the ME. Member genes that are *cis*-eGenes for these eSNPs are highlighted in orange.

To identify genetic regulators of the transcriptomic programmes captured by these modules, we tested for association between each module eigengene and the lead sepsis cis eSNPs identified here with >3 minor allele homozygotes in the cohort (12,335 SNPs). We found 241 significant associations for 30 modules across 32 loci (p<0.05/(12,335×106)=3.82e-8), which we termed module QTL (modQTL) (Supplementary Table 19). We then sought to replicate these modQTL using our previously published microarray gene expression data^7,12,13^ (n=135 samples overlapping RNA-seq, n=506 non-overlapping). We computed eigengene values for each module gene set with at least 5 genes measured on the microarray, and found that 16/29 lead modQTL replicated in non-overlapping patients (Supplementary Figure 17, Supplementary Table 20). We noted that replicating modQTL had greater correlation between the RNA-seq and microarray eigengene values computed for the samples included in both datasets (Supplementary Figure 18).

We noted that for the majority of modQTL (30/32) the module contained at least one *cis*-eGene for the modQTL SNPs. Whilst this could provide support for trans regulation of the rest of the module genes via a *cis*-eGene, the module QTL could also be a result of the *cis*-eGene(s) dominating the module eigengene and thus driving the overall modQTL signal. Moreover, several modQTL involved clusters of genes close to the SNP, with few or no distal genes included in the module. Such modQTL could be driven by a single eSNP regulating multiple genes in *cis*, or haplotypes comprising multiple *cis*-eSNPs with the inferred co-regulation driven by linkage disequilibrium between *cis*-regulatory elements for multiple nearby genes. We therefore performed a stringent sensitivity analysis to prioritise modQTL with evidence of trans regulation. We recalculated the module eigengenes excluding all *cis*-eGenes associated with the modQTL SNPs, and retested for association with the lead module SNP. This identified 7/32 robust modQTL that remained significant using the reduced module gene sets (Supplementary Figure 19, Supplementary Table 20). We then tested for mediation^54^ by the *cis*-eGene(s) of associations between the lead modQTL eSNP and the recalculated eigengene, and for all 7 modQTL found significant evidence of mediation by an implicated *cis*-eGene (Supplementary Table 21).

Several of these prioritised modQTL involved disease-relevant modules, where the module eigengene was also associated with clinical phenotypes (Figure 4b, Supplementary Table 14), highlighting regulators that may be contributing to disease heterogeneity through modulation of a gene network. For example, we identified a replicating modQTL for Module 92 (Figure 4c), which was associated with SRS and diagnosis and enriched for mature neutrophil marker genes^14,55^ (Figure 4a). It also included several HLA class I genes, for which we have increased confidence in the gene expression quantification due to incorporation of personalised HLA references in our RNA-seq mapping pipeline (Figure 4d). Of the two implicated cis-eGenes, *NLRC5,* a known regulator of HLA class I genes with a key role in inflammatory processes^56^, was significantly upregulated in SRS1 and was a significant mediator of the modQTL effect (Supplementary Figure 20).

We additionally identified and replicated a modQTL for module 47 (Figure 4e), which was associated with SRS1 status and mortality. The lead modQTL SNP, rs16843927, has been previously associated with inflammatory bowel disease through GWAS^33,34,57^ and we found evidence for colocalisation^39^ of the modQTL and GWAS signals (PP4=99.1%). The module primarily consists of a cluster of zinc finger genes on chromosome 19 involved in retroviral repression and was enriched for targets of STAT1 and ZNF274, with ZNF274 itself included in the module gene set (Figure 4f). The *cis*-eGenes for the modQTL SNPs, *SENP7* and *IMPG2*, were not module members, but we observed significant partial mediation (p=0.018) by *SENP7* of the association between the SNP and module gene expression (Supplementary Figure 20). *SENP7* has previously been reported to regulate many of the module 47 zinc finger genes in trans^55^, further supporting this association.

## Discussion

There are currently no immunomodulatory drugs available to treat the dysregulated immune response underlying sepsis, despite a large number of initially promising candidates^5^. Drug targets with genetic evidence are at least twice as likely to be successful in clinical trials^58,59^, so leveraging genetic associations for sepsis phenotypes is an important avenue to explore. Here, we have investigated the role of genetic variation in the sepsis response, finding evidence for genetic risk factors, and identified putative driver TFs and regulatory networks underlying inter-individual variation in the response to infection.

### Employing the GWAS approach in sepsis

We focused our GWAS on a composite molecular phenotype rather than an outcome such as mortality to improve power to detect genetic associations and aid mechanistic interpretation. While the lack of available expression data to assign SRS in external cohorts limits our ability to validate our findings, we did find one previously reported mortality-associated variant, rs72998754^60^, was in strong LD (r^2^=0.98 in 1KGP EUR) with nominally significant SNPs in our SRS GWAS (p=0.00015). Furthermore, through integrating our sepsis eQTL data, we find several plausible drivers of variation in the individual sepsis response. For example, one locus involves an eSNP for *NR2F6*, a transcription factor that represses transcription of key cytokines in CD4^+^ and CD8^+^ effector T cells including IL-2, IFNg, and TNFa^61,62^. The eSNP is located within an ATF3 ChIP peak^63^ and therefore may be specifically relevant in the context of stress and immune regulation.

### Context-specificity of eQTL effects

It has been widely reported that, despite their presumed regulatory activity, GWAS associations only have limited overlap with eQTLs^64^. One possible explanation for this observation is that the majority of eQTL studies have been conducted on healthy individuals, and regulatory variants may only be active in disease-relevant conditions, such as following immune activation. As sepsis represents an extreme and systemic response to infection, our sepsis eQTL results may thus help interpretation of risk variants for a broad range of immune and inflammatory diseases. Moreover, elucidating how environmental context impacts regulatory associations may improve understanding of how genetic variants contribute to complex traits. Additionally, we found that nearly 2000 sepsis eQTL signals had a significant interaction with SRS, source of sepsis, and/or measured cell proportions.

### Cell proportion differences as drivers of eQTL interactions

Transcriptomic regulation is known to be highly variable across cell types^65–67^. The eQTLs detected in bulk gene expression from a mixed cell population could therefore be affected by varying proportions of different cell types across patients, as well as in comparison to health. Development of sepsis and sepsis severity are associated with profound changes in leukocyte proportions; in particular, expansion of the neutrophil compartment has been observed in sepsis^6814,69^, COVID-19^70^, and ARDS^71^. This involves both mature neutrophils, known to be increased in sepsis, and immature neutrophil subpopulations that are rare or absent in the circulation of healthy individuals but are mobilised in response to infection. It is likely that some sepsis-enhanced or specific eQTLs derive from these leukocyte subpopulations. As immunosuppressive neutrophil subsets are associated with secondary infections in sepsis^72^, such effects could be of particular relevance to outcomes. However, around half our SRS1 magnifying interactions did not overlap with a cell type interaction and therefore differences in cell types do not explain all the effects observed, and additionally may not pinpoint the causative drivers involved.

### Identifying putative drivers of SRS

Heterogeneity across patients is a major obstacle to the identification of risk factors and the development of targeted treatments in sepsis^73,74^. Identification of the hub genes that modulate networks could enable personalised, targeted immunomodulation in sepsis. We identified 41 putative driver TFs with differential activity between SRS groups and for which the binding motif is also enriched in SRS interaction QTL.

The most significantly enriched motif was for the HIF-1 transcription factor complex comprising HIF1α and ARNT (HIF1β). We previously reported that a network centred on HIF1α was upregulated in the poor outcome SRS1 patients^7^, and we replicate this finding here, with both HIF1α and ARNT showing greater gene expression and elevated inferred activity in SRS1. Whilst HIF1α activation occurs in response to hypoxia and inflammatory signalling following infection, ARNT is constitutively expressed and its regulation is less well characterised^75,76^. HIF-1 has been widely identified as important in the sepsis response, inducing metabolic reprogramming in immune cells^77^ as well as affecting coagulation^78^, cell proliferation and apoptosis^79^, with a more pronounced impact in non-survivors^80,81^. The HIF1A-ARNT motif was also enriched among neutrophil interaction eQTLs, and we found HIF1α activity was correlated with the estimated proportions of platelets and immature neutrophils, and ARNT was correlated with cycling neutrophil progenitors and IL1R2+ immature neutrophils. When we previously characterised neutrophil subtypes in sepsis with an independent single cell dataset, we defined an IL1R2+ gene expression programme correlated with SRSq that was enriched for glycolytic and hypoxia-related pathways, highlighting a specific cell state where alterations in metabolism may be particularly important^14^.

The prioritised drivers also included CEBPA and CEBPB, regulators of steady-state and emergency granulopoiesis (EG) respectively^82^. Whilst the binding motifs tested are very similar, we have previously identified CEBPB in particular as a key factor in the SRS1 response in our single cell study^14^ and here we find a bigger difference in activity between SRS groups for CEBPB, further supporting the importance of EG in SRS1 with an independent approach. Furthermore, the inferred activity is correlated with immature neutrophil subsets as previously described^14^.

### Module QTL to identify genome-wide regulatory networks

Our eQTL interaction framework is one approach for identifying upstream regulators of groups of genes. We also utilised co-expression modules as a way of identifying putative trans regulatory networks with relevance to sepsis pathophysiology^83–85^. This approach identified a small number of associations, largely involving smaller modules (under 40 genes). Recent work has demonstrated that eQTL genes are less connected in co-expression networks compared to nearest genes of matched SNPs and GWAS associations (Mostafavi et al. 2022). Genes that are well-connected in gene regulatory networks are more important and thus require tightly controlled expression, resulting in higher constraint and fewer eQTLs due to negative selection. This reduces the power to discover cis-eQTLs for genes involved in large regulatory networks. Thus, our module analysis was likely powered to detect modQTL for genes involved in smaller regulatory networks.

Although modules in bulk expression cannot capture the cell type specificity of gene regulatory networks in a heterogeneous tissue like blood^86^, we were able to compare our gene sets with marker genes derived from single cell RNA sequencing in a smaller subset of individuals with sepsis^14^. This revealed that some of our modules captured cell-type signatures that are specific to immune cells in sepsis. Beyond the ability to generate data at scale, an advantage of identifying these signatures in bulk data is that they are more amenable to translation into a clinical setting. Of the 32 modQTL that we identified, module 47 was particularly interesting, with associations with SRS and mortality and module members including zinc finger genes involved in retroviral repression. We have previously described increased EBV viral load in SRS1 patients^87^. It is still unclear whether this association is a pathological effect rather than simply an epiphenomenon but this additional understanding of gene regulation indicates that further exploration is warranted.

### Limitations

There are necessarily limitations to our study. We adopted this eQTL framework because of the practical limitations of collecting sufficient samples in critical illness for well powered trans eQTL and GWAS, particularly given that SRS status requires the quantification of gene expression. We generated bulk RNA-seq to achieve sufficient sample sizes for eQTL mapping, but interpretation of cell type specific effects is limited to those that can be deconvoluted using a single cell reference. Given the importance of neutrophils in the response to sepsis, it is key to generate single cell data from whole blood rather than peripheral blood mononuclear cells; however it is still challenging to do this at scale. Technological advances will allow investigation of the more granular cell types contributing to the observed heterogeneity. The lack of availability of both genotyping and gene expression on the same sepsis patients also precluded validation in external cohorts. Our cohort is restricted to the two main sources of sepsis in the UK (CAP and FP). Whilst we observed a small number of eQTL effects differing by source of infection, potentially driven by location and/or pathogen type, we were not able to investigate pathogen specific effects due to insufficient microbiological information. Our cohort is of predominantly European ancestry and therefore population specific effects will not be identified. The ideal cohort design to investigate sepsis dependent eQTL effects would include pre-sepsis samples from the same individuals to allow a within-study interaction model. Given this is not feasible, we leveraged publicly available summary statistics^40,41^ to compare effect sizes^42^ between sepsis and health. However, technical differences between studies, such as cohort ancestry, sample size, experimental platform, the set of variants assessed and the availability of summary statistics can confound this comparison. We therefore minimised the impact of these variables by matching them to our cohort as closely as possible. Finally, there is currently very limited *in vivo* binding data on different transcription factors in primary cells, particularly in the disease context, and any single TF may have both activating and repressive effects across different genes and contexts. We have therefore used curated regulon and predicted binding sites to prioritise candidate regulatory drivers that could be suitable for longer term functional follow up.

## Conclusion

In conclusion, our eQTL interaction approach has identified factors putatively linking host genetic variation, cell subtypes, and the individual transcriptomic response to infection. Understanding the regulatory networks underlying patient heterogeneity could inform the development of immunomodulatory treatments and personalised medicine in sepsis.

## Methods

### Data generation and processing

#### Cohort description

We recruited adult patients (>18 years old) to the UK Genomic Advances in Sepsis (GAinS) study (NCT00121196) from 34 intensive care units (ICUs) between 16/11/2005 and 30/05/2018^7^. Ethics approval was granted nationally and locally for individual participating centres, and we obtained informed consent from the patient or their legal representative. Inclusion criteria were sepsis diagnosed according to ACCP/SCCM guidelines due to community acquired pneumonia (CAP) or faecal peritonitis (FP). CAP was defined as febrile illness associated with cough, sputum production, breathlessness, leukocytosis and radiological features of pneumonia acquired in the community or within two days of admission to hospital. FP was defined as inflammation of the peritoneal membrane secondary to faecal contamination, diagnosed by laparotomy. Exclusion criteria were immunosuppression, admission for palliative care only, and pregnancy.

#### Sample collection and processing

We extracted DNA from buffy coat or whole blood samples using the Qiagen DNA extraction protocol, the automated Maxwell Blood purification kit (Promega), or the QIAamp Blood Midi kit protocol (Qiagen). We took serial samples for RNA extraction on the first, third, and/or fifth day after ICU admission. We isolated the total leukocyte population at the bedside using the LeukoLOCK filter system (Life Technologies), and extracted purified RNA using the Total RNA Isolation Protocol.

#### Genotyping data generation and processing

We had previously generated genotyping data for 295 CAP patients and 730,525 SNPs using the Illumina HumanOmniExpress BeadChip^7^. We genotyped a further 655 patients at the Wellcome Sanger Institute using the Infinium CoreExome BeadChip (551, 839 SNPs), and 307 patients (including 38 repeat of samples that failed QC previously) at the Wellcome Centre for Human Genetics using the Infinium Global Screening Array BeadChip (654,027 SNPs).

We performed genotyping QC within each batch in Plink 1.9^88^ according to the methods described in Anderson et al 2010^89^. We excluded samples on the basis of discordant sex information, proportion of missing genotypes > 0.02, outlying heterozygosity rate, identity by descent (Pi_hat >=0.1875), and detection of sample mix-ups through comparison to RNAseq on the same patient (details below)^90^. We excluded variants if they had a missing data proportion > 0.05, MAF<0.01, and Hardy-Weinburg equilibrium P<1×10^-5^.

#### Imputation

We imputed each of the three genotyping data sets using the Haplotype Reference Consortium (HRC) release 1.1 panel and the Sanger Imputation Service^91^, following checks for strand, alleles, position, ref/alt assignments, and frequency differences versus the HRC (http://www.well.ox.ac.uk/~wrayner/tools/). Genotypes were phased using Eagle2 (v2.0.5^92^) and imputed using Positional Burrows-Wheeler Transform (PBWT^92,93^). We removed SNPs with an imputation info score <0.9, combined the data sets, and converted positions to build 38 coordinates using liftOver^92–94^. Finally, we excluded individuals with missingness >2% and SNPs with missingness ≥2%. This resulted in a final data set of 12,412,067 SNPs and 1,168 individuals. We calculated genotyping PCs on this combined data set using Plink^88^ and SNPs with MAF >1%.

#### RNA-sequencing and data processing

As described in Cano-Gamez et al (2022)^13^, we prepared stranded cDNA libraries for 909 samples from 695 sepsis patients using NEB Ultra II LIbrary Prep kits (Illumina) with NEBNext Poly(A) mRNA Magnetic isolation, and sequenced them across 3 S4 flow cells using an Illumina Novaseq 6000 system. We obtained a median of 40M 100bp paired-end reads per sample. We aligned reads to the reference genome (GRCh38.99) using STAR v2.7.3a^95^ (ENCODE recommended parameters) and quantified gene expression using featureCounts v2.0.0^96^ (https://github.com/wtsi-hgi/nextflow-pipelines/tree/2e5ac3cee33ca2a1ced2943bb7e366a7771a4d3c).

#### HLA allele calling to construct personalised references

To improve accuracy of HLA gene expression quantification, we created a personalised HLA reference for each patient. We imputed two-field resolution HLA types with arcasHLA 0.2.0^97^ using the IMGT/HLA database v 3.42.0 for fourteen HLA genes (*HLA-A, -B, -C, -E, -F, - DMA, -DMB, -DPA1, -DPB1, -DQA1, -DQB1, -DRA, -DRB1,* and *-DRB3*) and HIBAG v.1.4^97,98^ using a pre-trained multi-ethnic model for seven HLA genes (*HLA-A, -B, -C, -DPB1, -DQA1, -DQB1, and -DRB1*), from RNA-seq and genotyping data respectively. We then assessed the concordance between the RNA-seq and genotyping-based predictions as follows; 0: discordant calling for both alleles of a gene, 1: concordant call for one allele of a gene, 2: concordant calling for both alleles of a gene. We found greater concordance for HLA Class I alleles than class II (95.4% vs 91.2%) with >=1 concordant calls. We used HLA alleles from arcasHLA in cases with discordant calling. For alleles with missing calls from arcasHLA due to low RNA-seq read depth, alleles predicted by HIBAG were used. These imputed alleles were then used to define personalised reference sequences for HLA gene expression re-quantification.

In the IMGT/HLA database, untranslated exons are often missing, and UTR annotation is either absent or variable across alleles. As gene expression would be underestimated if reads originating from these regions were not mapped, we completed the personalised gene references by extending both the 5’ and 3’ sequences of each allele from IMGT based on primary reference sequences. In addition, for HLA-DRB2/4/7/8, we included two alternative reference sequences (GL000255 and GL000256) as these genes are missing in the primary reference sequences. Finally, we adjusted gene coordinates to reflect these extended allele sequences.

#### HLA allele mapping with personalised references

For each sample, we extracted reads mapping to the MHC (chr6:28500000-33400000, GRC38) and unmapped reads from the initial genome-wide mapping (see above). We aligned the extracted reads to the personalised reference sequences using STAR v2.7.3a. Depending on the heterozygosity of each gene, this produced up to 28 aligned BAM files per sample. We proceeded to assign reads to genes, following the method used in AltHapAlignR^99^. We merged all alignments from a given sample, retaining only uniquely mapped reads from each mapping run (bam file). We required that both reads in a pair were aligned to the exon region with at least one base on the same allele. For reads aligned to multiple genes, we compared the editing distance (NM value) between the reads and reference sequences and assigned the reads to the gene with the lowest editing distance. We discarded reads with the same editing distance in different genes. As the reads inputted to this HLA re-mapping included reads initially mapped to any region in the MHC and not just to the 14 HLA genes with personalised references, we also used this approach to assign reads previously mapped to any other MHC gene and remapped here. For reads with multiple gene assignments, we reassigned each read to the gene with the lowest editing distance. Finally, gene counts were recalculated for all MHC region genes.

#### Gene expression QC

We used MBV from QTLTools^90^ to identify mismatches between the genotyping and RNA-seq gene expression data. A mismatch, with or without sex mismatch in the genotyping or gene expression compared to the clinical information, could indicate a sample mixup. If we could determine the true sample identity, these were resolved; otherwise we excluded the affected patients from the analysis.

We excluded 45 samples following QC including mapping rate, PCA outliers, resolution of sample mix-ups and detection of contamination using VerifyBamID^100^. This resulted in a data set of 864 samples from 667 patients, including 135 samples repeated from previously published microarray data^7^. After targeted remapping of MHC region genes (see above), we filtered out features that did not have at least 10 reads in at least 5% of samples, retaining 20,412 genes for downstream analysis. We then normalised the data, using the trimmed mean of M-values method^101^ to estimate size factors for calculating counts per million, and added a pseudocount of 1 for log-transformation.

### GWAS

#### SRS assignment

All patient gene expression samples had Sepsis Response Signature (SRS) assignments from the SepstratifieR R package as described in Cano-Gamez et al^13^. We filtered our genotyping cohort to 997 patients with SRS assignments on at least 1 time point from any of microarray, RNA-seq, and qPCR. Where SRS1 status was available from multiple assays on the same time point, we found discrepant assignments for 10 samples. In these cases, we used SRS1 status from RNA-seq preferentially followed by microarray, resulting in 516 SRS1 samples and 818 non-SRS1 samples. We categorised patients as “Ever SRS1” if one or more time points (n1=736 patients, n2=185, n3=76) were assigned to SRS1. Our final cohort comprised 440 ever SRS1 patients and 557 never SRS1.

#### Heritability estimation

We used the GREML approach from GCTA^13,32,102,103^ to estimate the proportion of variation in the “ever SRS1” phenotype explained by common variants (MAF>1%). We used LD-pruned imputed autosomal genotype data (Plink^88^; --indep-pairwise 50 5 0.2) to calculate the GRM, and included age, age^2^, sex, and 7 genotyping principal components (PCs) as covariates in the model as fixed effects.

#### Genome-wide association study

We performed a GWAS for SRS1 ever vs never using Plink^88,104^, including age, age^2^, sex, and 7 genotyping principal components (PCs) as covariates in a logistic regression model. Independent loci were defined by clumping (r^2^<0.5). The coloc R package^39^ was used to perform colocalisation between GWAS and independent eQTL signals using default priors.

### eQTLs

#### cis-eQTL mapping

We restricted the genotyping data set to the 638 patients with RNA-seq gene expression data available, and to biallelic variants with MAF≥1% in this subset of individuals. Genotypes were coded as 0, 1, or 2 according to the number of copies of the minor allele carried by each patient. We filtered the RNA-seq data set to the 823 samples from 638 patients with genotyping data available. We calculated 30 PEER (probabilistic estimation of expression residuals) factors^105,106^ on the reduced gene expression dataset used for eQTL mapping, holding out rank-based inverse normal transformed (INT) clinically measured cell proportions (neutrophils, monocytes, lymphocytes), SRS1 status, site of infection (CAP/FP), and 7 genotyping PCs. For 5 samples with missing cell proportion information, we used the median value for the cohort.

We tested each autosomal gene for associations with SNPs within 1Mb of their transcriptional start site using the lme4 R package^107^. We used a linear mixed model to test each SNP-gene pair for an additive effect of genotype, including a random intercept effect for individual so that up to three serial samples per patient might be included to increase power^28^. We included the first seven principal components (PCs) of the genotyping data and the first 20 PEER factors of the gene expression data in the model to correct for systematic effects. This was based on including increasing numbers of PEER factors in the model to determine the “elbow” where adding additional factors did not result in a large increase in detected eQTLs (Supplementary Figure 3). We also included the covariates held out from the PEER factor calculation in the model.

We identified genes with a significant eQTL using a hierarchical approach for multiple testing correction^108^. Nominal p-values for the effect of the variant were obtained from a likelihood ratio test, using the anova function implemented in the lme4 package^107^. We used eigenMT^109^ to estimate the number of independent tests performed per gene, and from this calculated locally adjusted p-values for the peak association for each gene by Bonferroni correction. We then performed global p-value adjustment across the peak associations for all genes with the Benjamini-Hochberg method, and used a false discovery rate (FDR) threshold of 0.05 to identify genes with at least one significant eQTL (eGenes). We defined additional significant SNP-gene pairs for these eGenes as those with a nominal p-value less than the per-gene p-value threshold corresponding to the global FDR threshold of 0.05.

#### Conditional analysis

We then defined conditionally independent associations for each significant eGene using forward stepwise regression and backwards selection^36^. First, we repeated the *cis*-eQTL mapping, adjusting for the most significant SNP identified in the first round and using the gene-level significance threshold to identify independent secondary *cis*-eQTL signals. We repeated this iteratively, accounting for all the lead variants discovered in previous iterations to learn the number of independent signals. Once no new variants were identified, we terminated the forward stage and performed a backwards selection step, in which we re-tested each signal discovered in the forward step in turn. We included all other forward pass variants as covariates in the eQTL scan, and retained the most significant variant as the lead association for each independent signal. If we did not detect any significantly associated variants for a given signal in the backwards step, then the signal was dropped. Finally, we calculated effect sizes for each independent signal by fitting a model including all significant signal SNPs.

#### Overlap with external GWAS results

We queried the Open Target Genetics portal^33,34^ to look up previously reported associations for immune/inflammatory diseases and blood cell traits for SNPs of interest. We also obtained a list of SNPs associated with mortality in sepsis from Le et al 2019^110^ and Hernandez-Beeftink et al 2022^26^ to compare to our SRS GWAS results.

#### Interaction analysis

We tested the signal variants from the conditional eQTL analysis for interactions with SRS1 status, diagnosis (CAP vs FP) and cell proportion (monocyte, lymphocyte, or neutrophil), whilst also correcting for the effect of any independent signal variants. SNPs were only tested if there were at least two minor allele homozygote individuals in each subgroup or each half of all cell proportion distributions being tested. Interaction p values were calculated using a likelihood ratio test and corrected for multiple testing using the Benjamini-Hochberg method, and significant interactions identified using an FDR threshold of 0.05. To assess whether we had detected more interaction effects than would be expected by chance, we permuted SRS status across all samples or diagnosis across patients and repeated the interaction analysis 1000 times. Plots were made using the interactions R package.

#### Definition of sepsis-dependent eQTLs

We accessed the GTEx v8 whole blood summary statistics for European individuals through the GTEx Portal^40^, and the eQTLgen cis eQTL summary statistics from the eQTLgen Consortium website^41^. We compared z scores and significance for the lead SNPs for eGenes identified in GAinS, where the same SNP-gene pair was tested in the public dataset and the assessed allele could be disambiguated from strand designation. Significance in GTEx was defined as the gene having a qvalue < 0.05 in the *cis*-eGenes output, and the nominal p-value for the SNP being below the nominal p-value threshold for that gene.

We also used mashr^42^ to compare beta values between GAinS eGenes and GTEx, following the approach outlined in the mashr eQTL analysis vignette. Mashr (Multivariate Adaptive Shrinkage in R) compares effects on the same SNP-gene pairs across multiple conditions, employing empirical Bayes methods to estimate patterns of similarity. We used GTEx^40^ for this analysis, due to the greater technical similarity to GAinS and availability of beta values and standard errors. Briefly, we used a random 5% subset of the SNP-gene pairs tested in both GAinS and GTEx (“random” tests) to learn the correlation structure across null tests and the lead SNPs for sepsis eGenes (“strong” tests) to learn data-driven covariance matrices. We fitted the mashr model to the “random” tests, then used it to calculate posterior summaries on the “strong” tests from which to identify differing effects across conditions. We categorised the eQTLs significant in sepsis (mashr lfsr<0.05, n=8,122) as “shared” if the mashr posterior effect size was in the same direction as and within a factor of 0.5 of the GTEx effect size, or “context-dependent” if this was not the case. We classed those significant in both GAinS and GTex but with opposite directions of effects as “opposite effect” (n=53), and divided the remainder into those with bigger effects and/or only significant in sepsis (“sepsis-enhanced”, n=854, 10%) or in HV (“sepsis-dampened”, n=1,272)

#### Enrichment and network analysis

We used the XGR R package^111^ and Reactome annotations for pathway analysis, using Fisher’s Exact Test with the background defined as all genes tested in a given analysis. For example, when testing eGenes with an SRS interaction for enriched pathways, the background was all eGenes tested for an SRS interaction effect. P-values were adjusted using the Benjamini-Hochberg FDR procedure and an FDR threshold of 0.05 used to determine significance. We used a comparable approach to test for enrichment of gene sets defined in our results (e.g. sepsis-enhanced eGenes vs different types of interactions). We also used XGR to test for enrichment of interaction eSNPs vs non-interaction eSNPs in Roadmap Epigenomics Core 15-state Genome Segmentation annotations^46^ for primary peripheral immune cells with a one-tailed binomial test, and for exploration of subnetworks within eGenes with interactions using default parameters.

#### Transcription factor enrichment analysis

We used the SNP2TFBS^47,48^ database to identify instances of sepsis eSNPs introducing or interrupting predicted transcription factor binding sites. As the lead eSNP is not necessarily the causal variant, we first expanded the query SNPs to all SNPs in LD (r^2^≥0.8) with the signal SNPs in our cohort. We restricted the TF motifs considered to those annotated by Jaspar 2014 (JASPAR2014 R package^112^) as being defined in Homo sapiens (n=127/205 included in SNP2TFBS). We then collapsed the results for each independent eQTL signal, scoring each motif as having ≥1 or 0 binding sites altered by the signal SNP or its LD proxies. We then tested for enrichment of each TF motif with at least one SNP overlap, amongst eQTLs with a significant interaction effect compared to eQTLs with no significant interaction using a one-sided Fisher’s exact test. We adjusted the p-values obtained across all motifs using the Benjamini-Hochberg method. We permuted eGene interaction status 1000 times and repeated the enrichment analysis to determine how many enriched motifs we would expect to see by chance.

We inferred transcription factor activity from our RNA-seq dataset using the DoRothEA regulons^49^, restricted to human TFs with evidence levels A-C and a minimum regulon size of 5. We calculated a consensus activity score as implemented in the decoupleR R package^50^. We tested for differential activity between SRS groups using a linear mixed model with a random intercept for individual, adjusting the p-values for multiple testing using the Benjamini-Hochberg method. We permuted SRS status and repeated the differential activity analysis to determine how many TFs we would expect to have differential activity by chance.

We matched the transcription factors for which we had inferred activity to the motifs assessed by SNP2TFBS^47^ to pinpoint TFs with evidence of relevance to SRS in both analyses. Using the first available sample for each patient, we used Spearman correlation to identify relationships between TF activity and cell proportions estimated using CIBERSORTx^113^ based on a sepsis single cell reference set^14^.

### Co-expression modules

#### Co-expression module discovery

We identified co-expression modules in the gene expression data using the WGCNA R package^52^ as follows. First, to control for technical variation during module identification^114^, we regressed the top 20 gene expression PCs out from the logCPM gene expression matrix. So that we only considered between-individual correlation, we replaced the gene expression value in each sample from the same individual with their mean gene expression^115^. We then calculated the biweight midcorrelation matrix for the residual gene expression to generate a similarity matrix using the bicor function from the WGCNA R package^52^. We used spatial quantile normalisation implemented in the spqn R package^116^ to account for the mean-correlation bias in the similarity matrix, and applied the normalize_correlation function to the similarity matrix with 21 blocks of size 1000 and block 18 as the reference group.

We determined a soft threshold value of 4 for the similarity matrix using the pickSoftThreshold function. We used this soft threshold to build an unsigned adjacency matrix using the adjacency function, which we used in turn to calculate the topological overlap metric (TOM) matrix using the TOMsimilarity function. We applied the dynamic tree cut algorithm included in the WGCNA package as the cutreeDynamic function to generate modules with default parameters and a minimum cluster size of 10. Similar modules were merged based on the similarity of their module eigengenes using the mergeCloseModules function with a cut height of 0.1. For a module, the eigengene was defined as the first PC of the gene expression data of the genes present in the module. We calculated the module eigengenes for the final set of modules using the moduleEigengenes function.

#### Module Annotation

We tested for enrichment of Reactome pathways among module member genes using XGR, and for downstream targets of different transcription factors using DoRothEA regulons^49^ and Fisher’s Exact Test. In each case, p-values were adjusted using Benjamini-Hochberg FDR correction and a threshold of 0.05 used. The set of expressed genes was used as the background for enrichment.

We used cell markers from the xCell R package^53^ and a single-cell RNA-seq sepsis dataset^14^ to identify cell-type-specific modules. The xCell signatures were derived based on differential gene expression from large transcriptomic studies of individual cell types and built to minimise classification error. Enrichment of gene signatures was performed using a hypergeometric test using the phyper function in R. The entire set of expressed genes was considered the background for enrichment. P-values were corrected using the Benjamini-Hochberg FDR procedure. Since multiple transcriptomic studies assayed the same cell types in xCell, one cell type often had multiple signatures. The median odds of enrichment per cell type in xCell were reported for any signatures that passed a q-value cutoff of 0.05. In contrast, each cell type in the Kwok et al. 2023 study had one set of markers. Signatures for cell types passing a q-value cutoff of 0.05 were reported. Enrichment was calculated as the ratio between the proportion of signature genes in the module and the proportion of signature genes in the entire set of expressed genes.

We tested for association between 28-day survival and each eigengene using a Cox proportional hazards model, as implemented in the survival R package using the coxph function. For each patient, the value of the eigengene at the last time point assayed was used as a predictor for the survival function. For all other patient phenotypes (cell proportions, SRS1 status, diagnosis and time point), we used a linear mixed model to test for differences in eigengene expression between groups. Measured cell proportions were inverse normal transformed and missing values replaced with the median value. P-values were corrected for multiple testing using the Benjamini-Hochberg procedure.

#### Module QTL

We tested the 12,335 unique signal eSNPs from our conditional sepsis eQTLs with >3 minor allele homozygotes for associations with all module eigengenes. As in the cis-eQTL analysis, we included seven genotyping PCs, 20 PEER factors, SRS status (SRS1 versus non-SRS1), diagnosis (CAP versus FP), and transformed cell proportions as fixed-effect covariates. A genome-wide threshold of 3.82×10^!”^ was used based on a Bonferroni correction accounting for the number of SNPs and number of modules tested. We defined loci for each module by constructing 1 Mb windows around each module QTL SNP and merging those that overlapped.

#### ModQTL replication

We sought to replicate the 32 lead modQTL associations using our previously published microarray gene expression data^13^. First, for each module gene set with at least 5 genes passing QC in the microarray (Modules 97, 88, 103 were not replicable), we computed eigengene values in the full microarray cohort (n=676 samples) using the svd R function. We assessed eigengene correlation across technologies based on 135 patient samples included in both the microarray and the RNA-seq cohort (Spearman’s rho). We confirmed that all lead modQTL SNPs had a MAF ≥1% in the non-overlapping microarray sample set with genotyping data (n=506 samples from 361 patients). We then tested for replication of the association of the module eigengenes with the lead modQTL SNP in these samples using the same model design as above (p-value threshold 0.05). We used the signs of the beta values and of the correlation coefficients from the 135 repeated samples to assess concordance of direction of effects between the RNA-seq and microarray cohorts.

#### ModQTL sensitivity analysis

We recalculated the module eigengenes in the RNA-seq cohort excluding all *cis*-eGenes for which any associated SNPs were lead eSNPs, and retested for association with the lead module SNP using the same model as above. Significance was determined using the same genome-wide threshold.

#### Mediation analysis

We then tested for mediation of associations between the lead modQTL eSNP (treatment) and the recalculated eigengene (outcome) by the modQTL SNPs’ target *cis*-eGene(s) (mediator, each tested separately) using the mediation R package^54^. All covariates used in the main modQTL model were included. A quasi-Bayesian approximation was used for confidence intervals with 1000 simulations, and effect sizes represent the effect of 1 additional copy of the minor allele.

## Supplementary Figures

**Figure S1: Genetic contribution to the SRS1 phenotype.**

The contribution of common SNPs to variation in SRS status, summarised for each patient as “Ever assigned to SRS1 in the first 5 days in ICU” vs “Never assigned to SRS1”, was estimated using GCTA. Estimates of variance in the phenotype (Vp), the variance explained by environmental factors (V(e)) and by common SNPs (V(G)) are shown as a forest plot. The proportion of the phenotypic variance explained by common SNPs collectively is stated underneath.

**Figure S2: Genome-wide association study for SRS1ever vs SRS1never.**

Common SNPs (MAF≥1%) were tested for association with the SRS1ever vs never phenotype. qqplot showing the observed p-value distribution plotted against the expected distribution.

**Figure S3: Impact of the number of PEER factors included in the eQTL model.**

As more PEER factors are included in the eQTL model, the number of eQTLs detected on chr1 increases non-linearly (*left*). The number of additional eQTLs detected with every additional 5 PEER factors decreases rapidly (*right*).

**Figure S4: Distribution of conditional lead eQTL SNPs around the eGene transcriptional start site (TSS).**

The base pair distance from the eSNP to its eGene’s TSS was calculated and plotted as density plots by the rank of the conditional eQTL signal. Primary signals (red) were closer to the TSS than secondary signals (green), which were in turn closer than tertiary and greater signals (blue).

**Figure S5: Colocalisation of an SRS GWAS signal with eQTL signals for *NR2F6* and *OCEL1*.**

A SNP that passed the genome-wide suggestive threshold in the SRS GWAS was also a significant eQTL for *NR2F6* and *OCEL1,* and testing for colocalisation indicated that the same causal SNP was driving the three signals. Each point is a variant, with significance for each association plotted against genomic location and colour indicating LD (r^2^) with the lead SNP from the GWAS association.

**Figure S6: Replication of eQTL results from a microarray sepsis cohort.**

Comparison of beta values for all SNP-gene pairs with nominal significance in our previous microarray eQTL study^7^ that were also tested in this RNA-seq cohort. Blue colour indicates significance in the current study.

**Figure S7: Distribution of source of sepsis interaction QTL from permutation analysis.**

Source of sepsis was permuted across individuals and the eQTL interaction analysis was repeated, with the number of significant interactions for each permutation plotted as a histogram. The observed result is marked with a red arrow.

**Figure S8: Comparison of sepsis eQTLs to GTEx and eQTLgen results.**

Comparison of z-scores for sepsis lead SNP-eGene pairs and GTEx version 8 whole blood eQTL results from European individuals (top) and eQTLGen (bottom).

**Figure S9: Characteristics of sepsis-enhanced eQTL variants.**

Density plots demonstrating how eSNPs involved in sepsis-enhanced eQTLs differ from eSNPs involved in eQTLs with comparable effect sizes to GTEx in terms of (left) MAF, (middle and right) distance to the transcriptional start site (TSS) of the eGene.

**Figure S10: Distribution of SRS interaction QTL from permutation analysis.**

SRS status was permuted across samples and the eQTL interaction analysis was repeated, with the number of significant interactions for each permutation plotted as a histogram. The observed result is marked with a red arrow.

**Figure S11: Cell proportion interaction QTL results.**

eQTL interactions with measured cell proportions. Each point represents an independent eSNP-eGene pair, with the interaction effect size plotted against the genotype effect. eQTLs with bigger effects with increased cell proportion are found in the top right and bottom left quadrants. Red colour indicates a significant interaction between genotype and cell proportion (FDR<0.05), with the most significant results labelled with the eGene name.

**Figure S12: Distribution of transcription factor binding site enrichment results from permutation analysis.**

eQTL interaction status was permuted across all eQTL signals and the TFBS enrichment tests were repeated, with the number of significantly enriched motifs for each permutation plotted as a histogram. The observed result is marked with a red arrow.

**Figure S13: Distribution of DoRothEA inferred TF activity differences from permutation analysis.**

SRS status was permuted across samples and inferred transcription factor activity compared between groups, with the number of significantly differing TFs for each permutation plotted as a histogram. The observed result is marked with a red arrow.

**Figure S14: Histogram of co-expression module size.**

The number of genes in each module ranged from 11 to 1,785.

**Figure S15: Module eigengene associations with sepsis endophenotypes.**

Heatmap showing significant associations between all module eigengenes (MEs) and clinical phenotypes. MEs were tested for differential expression with measured cell proportions, SRS1 status, diagnosis (CAP or FP) and time point (day 1, 3, 5) using a linear mixed model. Association of each ME with survival up to 28 days was tested using a Cox proportional hazards model.

**Figure S16: Enrichment of xCell marker genes in module members.**

Modules were tested for enrichment of xCell^53^ gene signatures derived from large whole blood transcriptomic studies. Modules shown had significant enrichment for at least one signature.

**Figure S17: Replication of modQTL effects in a microarray sepsis cohort.**

Forest plot of modQTL replicated in a validation cohort. Of the 29 modQTL that could be tested, 16 were replicated with consistent direction of effect. Effect sizes from the discovery

RNA-seq data set and the replication microarray data set are shown as points with 95% confidence intervals as lines.

**Figure S18: Correlation of module eigengenes across technologies by replication status.**

Module eigengenes were calculated using the same module gene sets in a microarray cohort with 135 overlapping samples. Similarity between module eigengenes were tested using Spearman’s rho for the overlapping samples. Module QTL that replicated with the non-overlapping samples had better correlated eigengenes between the two datasets.

**Figure S19: ModQTL sensitivity analysis.**

Module eigengenes were recalculated excluding any eGenes for the associated modQTL SNPs, and the modQTL retested. The new beta value is plotted against the original for the same SNP-ME pair, and coloured by whether the association remained significant.

**Figure S20: ModQTL mediation results.**

We tested for mediation of associations between the lead modQTL eSNP and the recalculated eigengene by the modQTL SNPs’ target *cis*-eGene(s). The Average Causal Mediated Effect (ACME), Average Direct Effect (ADE) and total effects are plotted for two modQTL of biological interest.

## Supplementary Tables

**Table S1:** SRS1ever vs never GWAS summary statistics

**Table S2:** SRS1ever vs never GWAS loci

**Table S3:** Sepsis *cis*-eQTLs (initial pass)

**Table S4:** Conditional sepsis *cis*-eQTLs

**Table S5:** Comparison with microarray eQTLs

**Table S6:** Source of sepsis eQTL interactions

**Table S7:** Sepsis dependence

**Table S8:** SRS eQTL interactions

**Table S9:** Cell proportion eQTL interactions

**Table S10:** SNP2TFBS motif enrichment in SRS interaction loci

**Table S11:** DoRothEA inferred TF activity differences by SRS

**Table S12:** Correlation between estimated cell proportions and TF activities

**Table S13:** Co-expression module membership

**Table S14:** Module endophenotype associations

**Table S15:** Module pathway enrichment

**Table S16:** Module xCell marker genes enrichment

**Table S17:** Module sepsis cell marker genes enrichment

**Table S18:** Module TF regulon enrichment

**Table S19:** modQTL

**Table S20:** modQTL replication in microarray cohort and sensitivity analysis results

**Table S21:** modQTL mediation results

## Acknowledgements

We thank all the patients, patient families, nurses, and clinicians who participated in the UK Genomic Advances in Sepsis (GAinS) study. We are grateful to Giuseppe Scozzafava for maintaining the sample biobank and to the Sanger Institute’s Scientific Operations team for generating the RNA-seq data. We thank Guillaume Noell and the Wellcome Sanger Institute’s Human Genetics Informatics (HGI) team for mapping the bulk RNA-sequencing reads. We are grateful to all members of the Davenport lab, and particularly to Alex Tokolyi for advice on the mediation approach. The Genotype-Tissue Expression (GTEx) Project was supported by the Common Fund of the Office of the Director of the National Institutes of Health, and by NCI, NHGRI, NHLBI, NIDA, NIMH, and NINDS. Data used for specific analyses described in this manuscript were obtained from the GTEx Portal on 23/10/19.

## Author contributions

E.E.D., K.L.B., and J.C.K. conceptualised the study. E.E.D. and J.C.K. supervised the study. Funding was acquired by J.C.K., E.E.D. and N.S. C.J.H., J.C.K., S.M. and the GAinS investigators recruited the patients. K.L.B., Y.M., C.G.G. and A.J.K. performed the experimental work. A.J.K., E.C-G. and S.M. provided resources. K.L.B., N.M. and W.L. performed the data analysis. K.L.B, E.E.D., N.M., W.L. and J.C.K. interpreted the results. K.L.B., E.E.D., N.M., and W.L drafted the manuscript. Review and editing was carried out by all authors who read, provided input on and approved the paper.

For the purpose of Open Access, the author has applied a CC BY public copyright licence to any Author Accepted Manuscript version arising from this submission.

## GAinS Investigators

In addition to the GAinS Investigators who are authors (K.L.B., Y.M., C.G.G., E.E.D., C.J.H., J.C.K., and S.M.), the following GAinS investigators, listed alphabetically by institution, were involved in patient recruitment, sample collection, or sample processing:

Nigel Webster^7^, Helen Galley^7^, Jane Taylor^7^, Sally Hall^7^, Jenni Addison^7^, Sian Roughton^7^, Heather Tennant^7^, Achyut Guleri^8^, Natalia Waddington^8^, Dilshan Arawwawala^9^, John Durcan^9^, Alasdair Short^9^, Karen Swan^9^, Sarah Williams^9^, Susan Smolen^9^, Christine Mitchell-Inwang^9^, Tony Gordon^10^, Emily Errington^10^, Maie Templeton^10^, Pyda Venatesh^11^, Geraldine Ward^11^, Marie McCauley^11^, Simon Baudouin^12,28^, Charley Higham^12^, Jasmeet Soar^13^, Sally Grier^13^, Elaine Hall^13^, Stephen Brett^14^, David Kitson^14^, Robert Wilson^14^, Laura Mountford^14^, Juan Moreno^14^, Peter Hall^15^, Jackie Hewlett^15^, Christopher Garrard^16^, Julian Millo^16^, Duncan Young^16^, Paula Hutton^16^, Penny Parsons^16^, Alex Smiths^16^, Roser Faras-Arraya^16^, Jasmeet Soar^17^, Parizade Raymode^17^, Jonathan Thompson^18^, Sarah Bowrey^18^, Sandra Kazembe^18^, Natalie Rich^18^, Prem Andreou^18^, Dawn Hales^18^, Emma Roberts^18^, Simon Fletcher^19^, Melissa Rosbergen^19^, Georgina Glister^19^, Jeronimo Moreno Cuesta^20^, Julian Bion^21^, Joanne Millar^21^, Elsa Jane Perry^21^, Heather Willis^21^, Natalie Mitchell^21^, Sebastian Ruel^21^, Ronald Carrera^21^, Jude Wilde^21^, Annette Nilson^21^, Sarah Lees^21^, Atul Kapila^22^, Nicola Jacques^22^, Jane Atkinson^22^, Abby Brown^22^, Heather Prowse^22^, Anton Krige^23^, Martin Bland^23^, Lynne Bullock^23^, Donna Harrison^23^, Gary Mills^24,25^, John Humphreys^24,25^, Kelsey Armitage^24,25^, Shond Laha^26^, Jacqueline Baldwin^26^, Angela Walsh^26^, Nicola Doherty^26^, Stephen Drage^27^, Laura Ortiz-Ruiz de Gordoa^27^, Sarah Lowes^27^, Charley Higham^28^, Helen Walsh^28^, Verity Calder^28^, Catherine Swan^28^, Heather Payne^28^, David Higgins^29^, Sarah Andrews^29^, Sarah Mappleback^29^, D Watson^30,31^, Eleanor McLees^30,31^, Alice Purdy^30,31^, Martin Stotz^32^, Adaeze Ochelli-Okpue^32^, Stephen Bonner^33^, Iain Whitehead^33^, Keith Hugil^33^, Victoria Goodridge^33^, Louisa Cawthor^33^, Martin Kuper^34^, Sheik Pahary^34^, Geoffrey Bellingan^35^, Richard Marshall^35^, Hugh Montgomery^35^, Jung Hyun Ryu^35^, Georgia Bercades^35^, Susan Boluda^35^, Andrew Bentley^36^, Katie Mccalman^36^, Fiona Jefferies^36^, Narelle Maugeri^3^, Jayachandran Radhakrishnan^3^ and Alice Allcock^3^.

^7^Aberdeen Royal Infirmary, Aberdeen AB25 2ZN, UK.

^8^Blackpool Victoria Hospital, Blackpool FY3 8NR, UK.

^9^Broomfield Hospital, Chelmsford CM1 7ET, UK.

^10^Charing Cross Hospital, London W6 8RF, UK.

^11^Coventry and Warwickshire University Hospital, Coventry CV2 2DX, UK.

^12^Freeman Hospital, Newcastle upon Tyne NE7 7DN, UK.

^13^Frenchay Hospital, Bristol, UK and Southmead Hospital, Bristol BS16 1JE, UK.

^14^Hammersmith Hospital, London W12 0HS, UK.

^15^Huddersfield Royal Infirmary, Huddersfield HD3 3EA, UK.

^16^John Radcliffe Hospital, Headington, Oxford OX3 9DU, UK.

^17^Kettering General Hospital, Kettering NN16 8UZ, UK.

^18^Leicester Royal Infirmary, Leicester LE1 5WW, UK.

^19^Norfolk and Norwich University Hospital, Norwich NR4 7UY, UK.

^20^North Middlesex Hospital, London N18 1QX, UK.

^21^Queen Elizabeth Hospital, Birmingham B15 2GW, UK.

^22^Royal Berkshire Hospital, Reading RG1 5AN, UK.

^23^Royal Blackburn Hospital, Blackburn BB2 3HH, UK.

^24^Royal Hallamshire Hospital, Sheffield S10 2JF, UK.

^25^Northern General Hospital, Sheffield S5 7AU, UK.

^26^Royal Preston Hospital, Preston PR2 9HT, UK.

^27^Royal Sussex County Hospital, Brighton BN2 5BE, UK.

^28^Royal Victoria Infirmary, Newcastle upon Tyne NE1 4LP, UK.

^29^Southend Hospital, Westcliff-on-Sea SS0 0RY, UK.

^30^St Bartholomew’s Hospital, London EC1A 7BE, UK.

^31^Royal London Hospital, London E1 1FR, UK.

^32^St Mary’s Hospital, London W2 1NY, UK.

^33^James Cook University Hospital, Middlesbrough TS4 3BW, UK.

^34^Whittington Hospital, London N19 5NF, UK.

^35^University College London Hospital, UCLH, London NW1 2BU, UK.

^36^Wythenshawe Hospital, Manchester M23 9LT, UK.

