## Supplementary Figures for "eQTLs identify regulatory networks and drivers of variation in the individual response to sepsis"

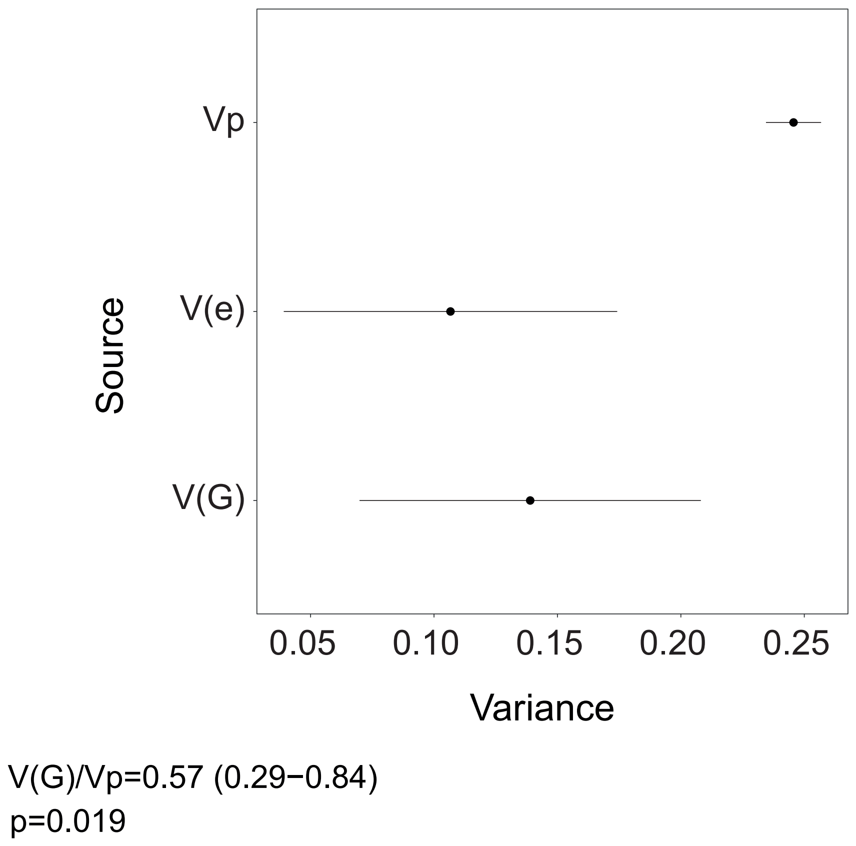


**Figure S1: Genetic contribution to the SRS1 phenotype.**
The contribution of common SNPs to variation in SRS status, summarised for each patient as “Ever assigned to SRS1 in the first 5 days in ICU” vs “Never assigned to SRS1”, was estimated using GCTA. Estimates of variance in the phenotype (Vp), the variance explained by environmental factors (V(e)) and by common SNPs (V(G)) are shown as a forest plot. The proportion of the phenotypic variance explained by common SNPs collectively is stated underneath.

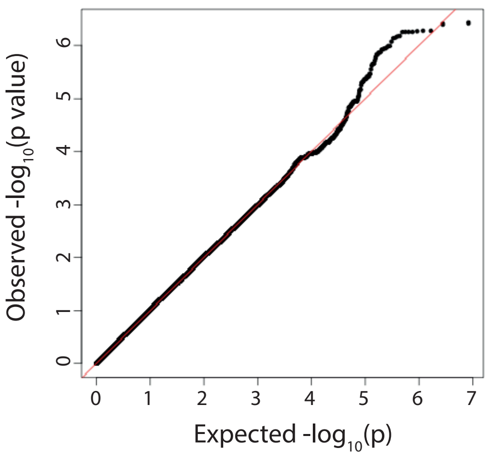


**Figure S2: Genome-wide association study for SRS1ever vs SRS1never.**
Common SNPs (MAF>1%) were tested for association with the SRS1ever vs never phenotype. qqplot showing the observed p-value distribution plotted against the expected distribution.

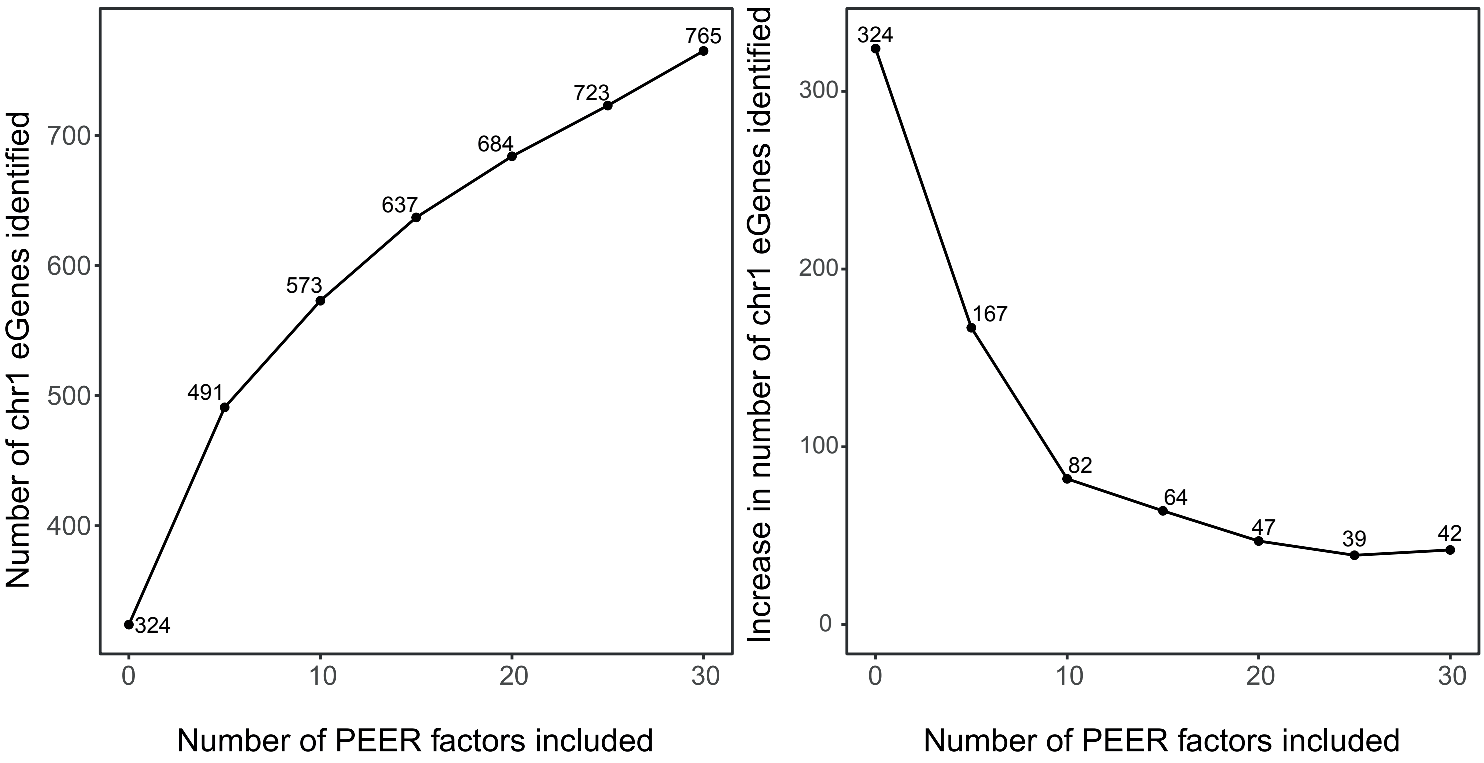


**Figure S3: Impact of the number of PEER factors included in the eQTL model.**
As more PEER factors are included in the eQTL model, the number of eQTLs detected on chr1 increases non-linearly (*left*). The number of additional eQTLs detected with every additional 5 PEER factors decreases rapidly (*right*).

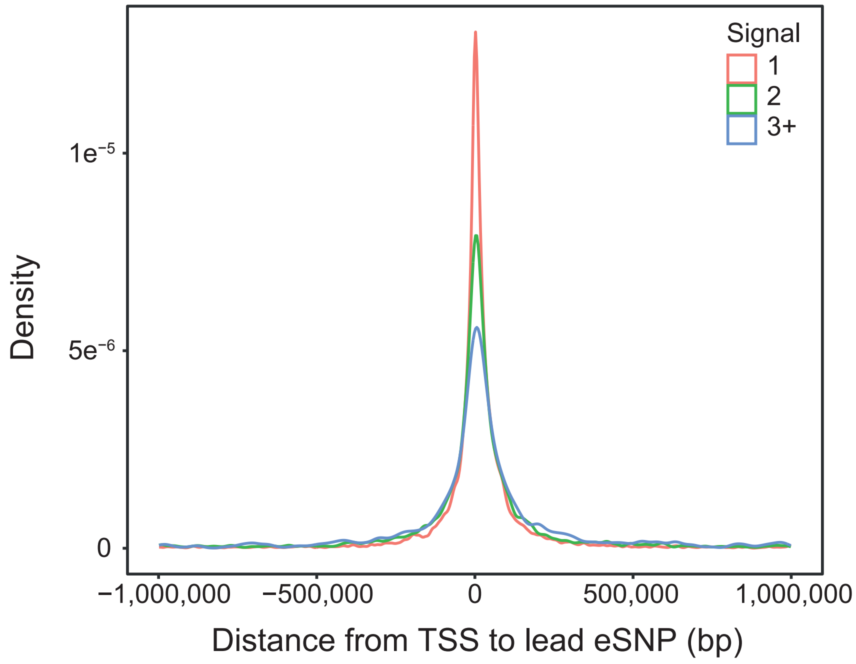


**Figure S4: Distribution of conditional lead eQTL SNPs around the eGene transcriptional start site (TSS).**
The base pair distance from the eSNP to its eGene's TSS was calculated and plotted as density plots by the rank of the conditional eQTL signal. Primary signals (red) were closer to the TSS than secondary signals (green), which were in turn closer than tertiary and greater signals (blue).

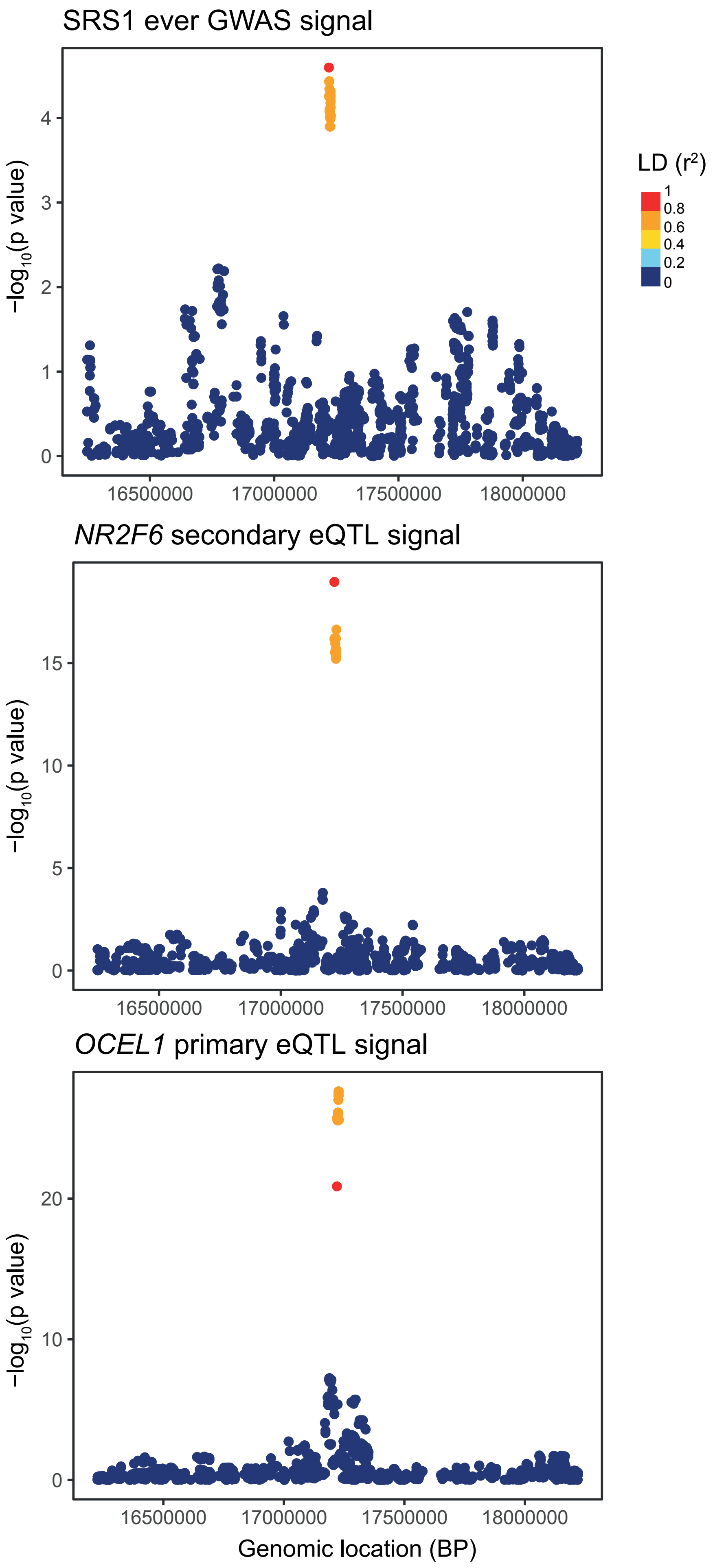


**Figure S5: Colocalisation of an SRS GWAS signal with eQTL signals for *NR2F6* and *OCEL1*.** A SNP that passed the genome-wide suggestive threshold in the SRS GWAS was also a significant eQTL for *NR2F6* and *OCEL1*, and testing for colocalisation indicated that the same causal SNP was driving the three signals. Each point is a variant, with significance for each association plotted against genomic location and colour indicating LD ($r^2$) with the lead SNP from the GWAS association.

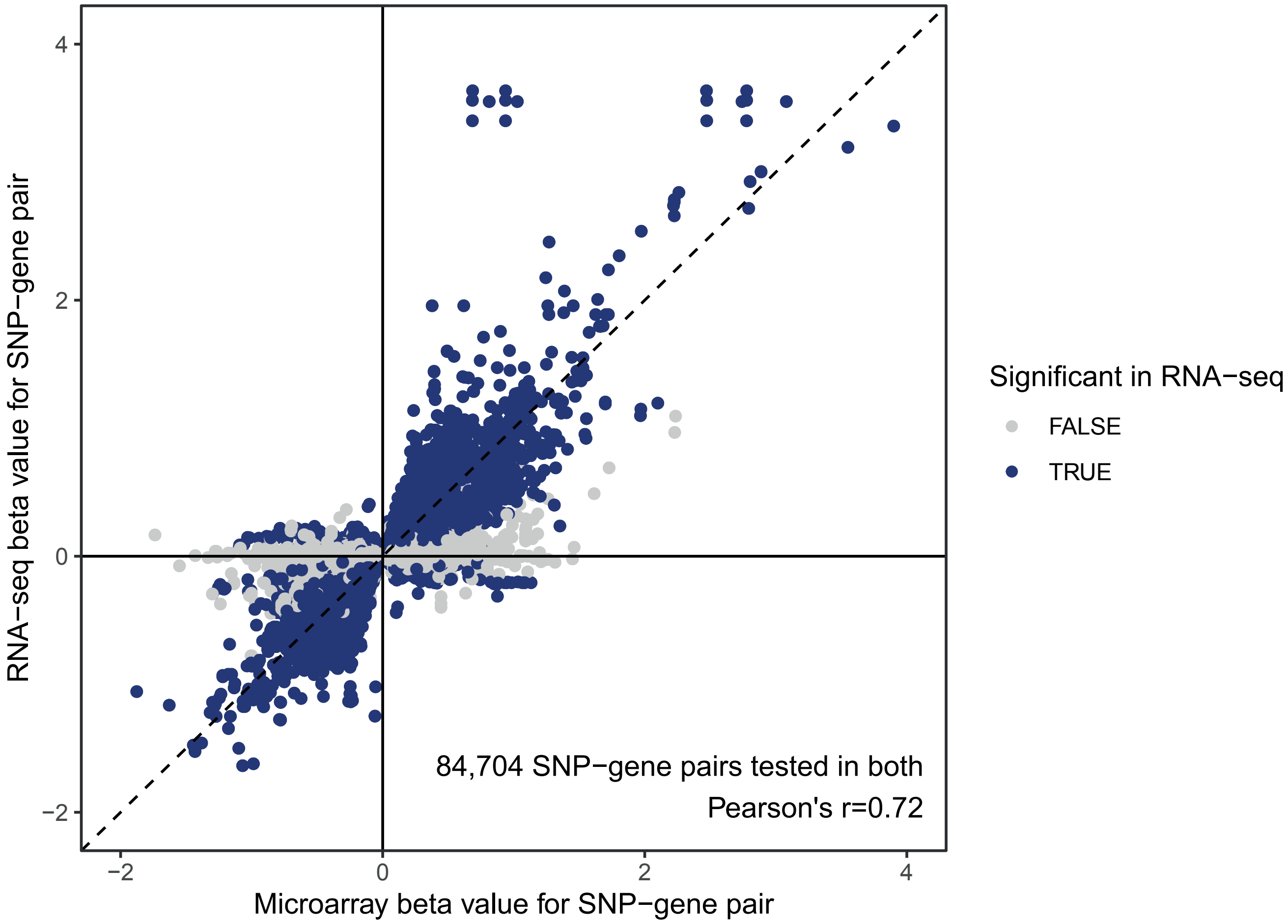


**Figure S6: Replication of eQTL results from a microarray sepsis cohort.**
Comparison of beta values for all SNP-gene pairs with nominal significance in our previous microarray eQTL study that were also tested in this RNA-seq cohort. Blue colour indicates significance in the current study.

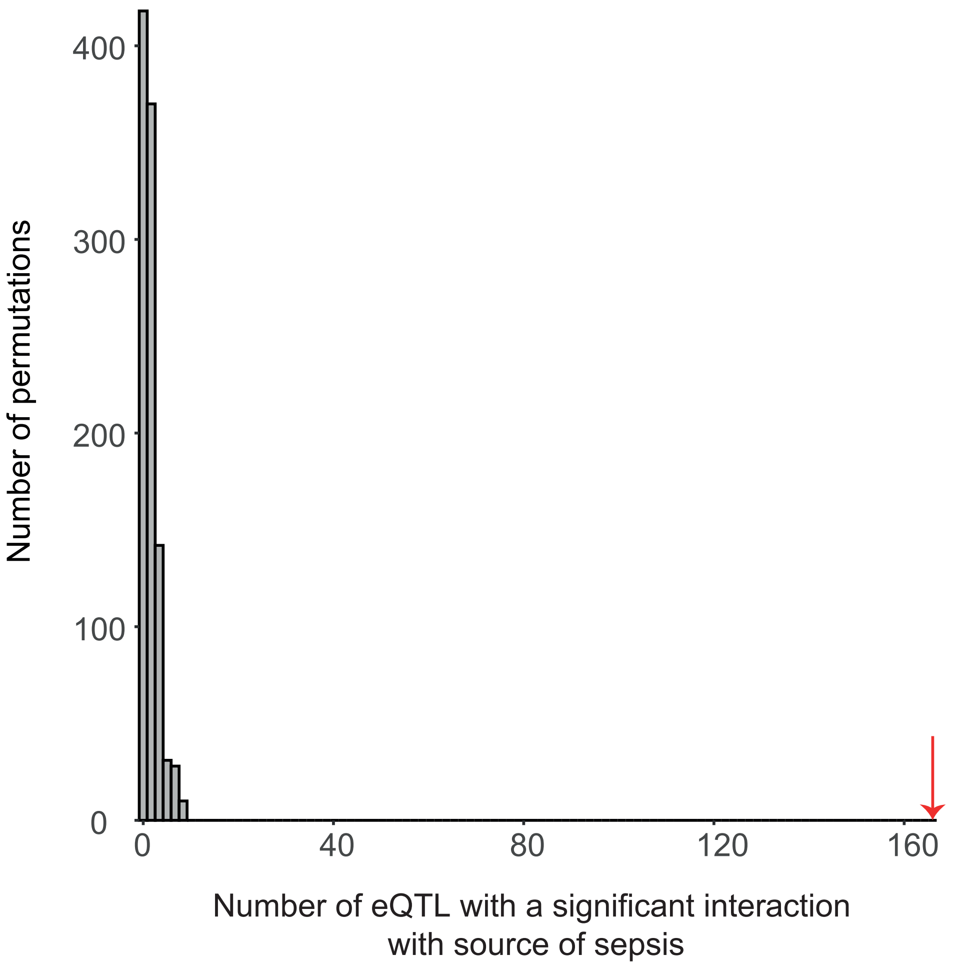


**Figure S7: Distribution of source of sepsis interaction QTL from permutation analysis.** Source of sepsis was permuted across individuals and the eQTL interaction analysis was repeated, with the number of significant interactions for each permutation plotted as a histogram. The observed result is marked with a red arrow.

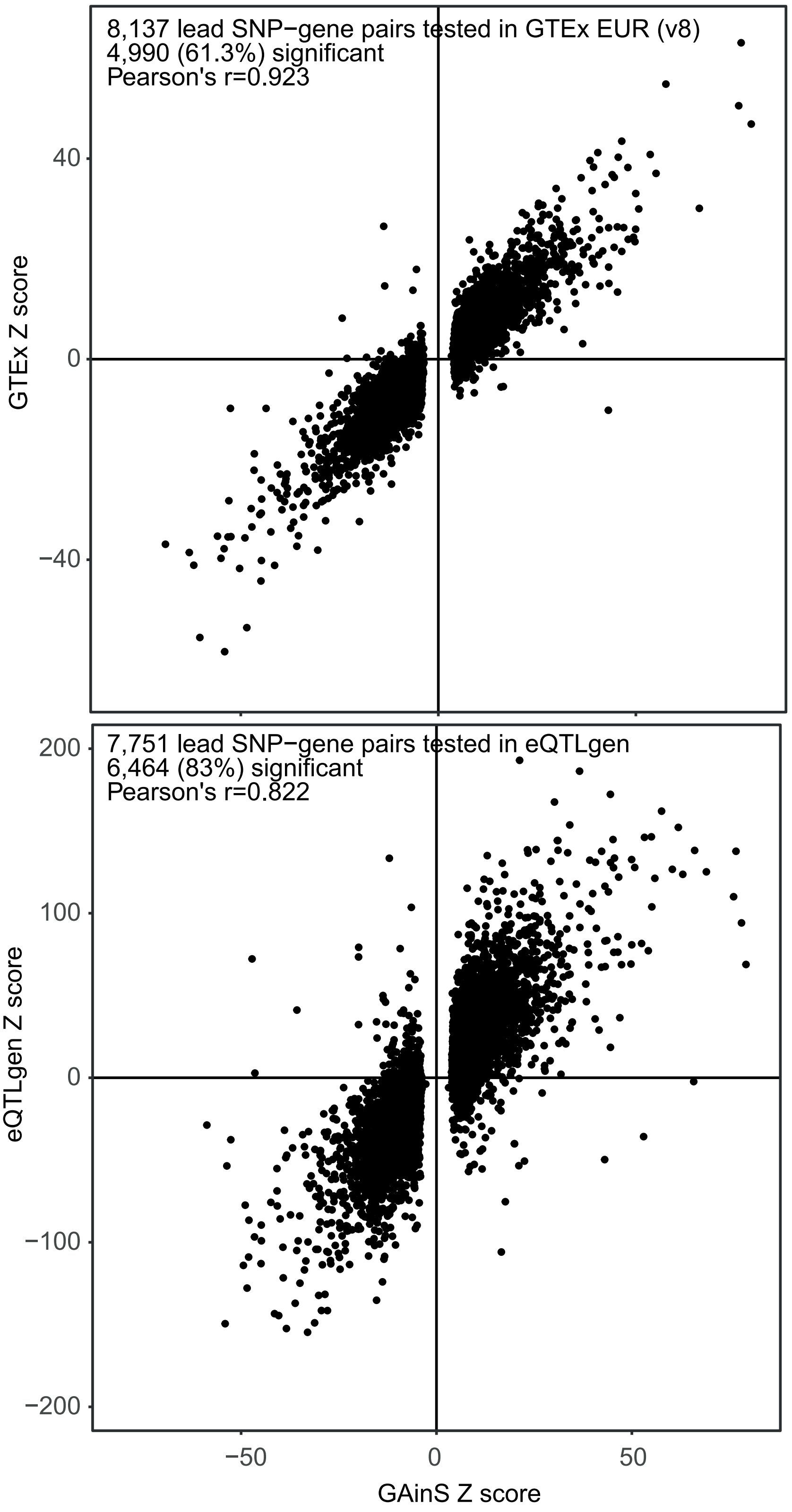


**Figure S8: Comparison of sepsis eQTLs to GTEx and eQTLgen results.**
Comparison of z-scores for sepsis lead SNP-eGene pairs and GTEx version 8 whole blood eQTL results from European individuals (top) and eQTLGen (bottom).

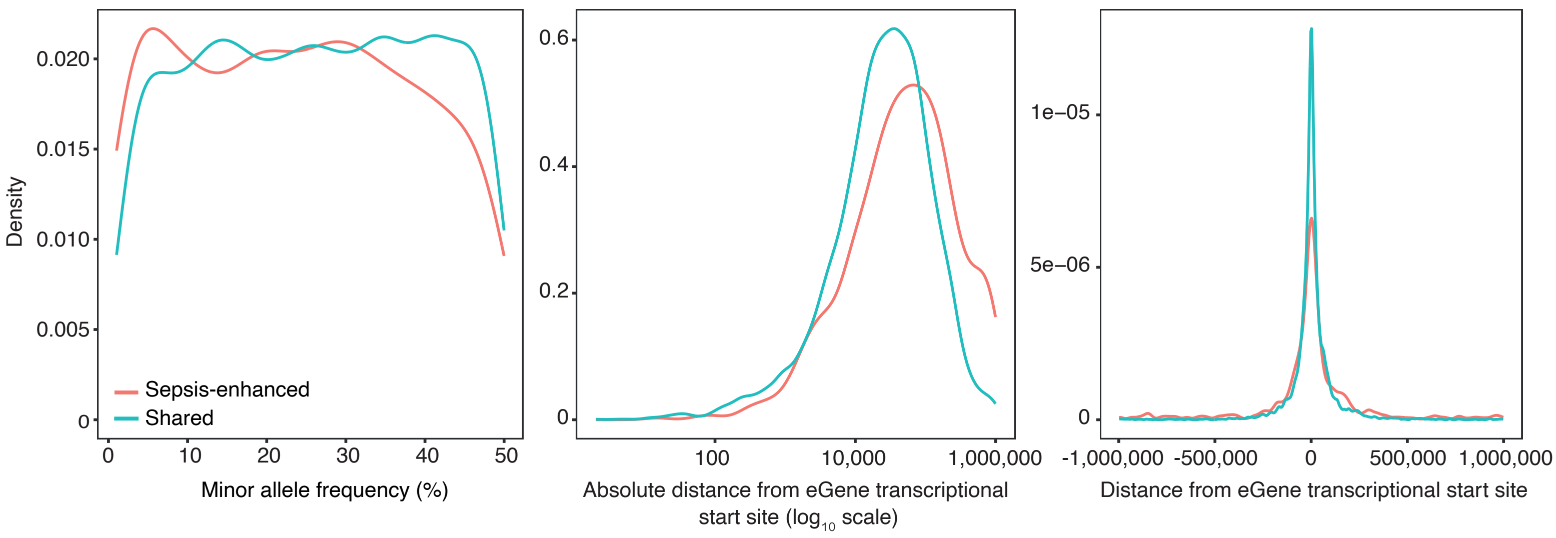


**Figure S9: Characteristics of sepsis-enhanced eQTL variants.**
Density plots demonstrating how eSNPs involved in sepsis-enhanced eQTLs differ from eSNPs involved in eQTLs with comparable effect sizes to GTEx in terms of (left) MAF, (middle and right) distance to the transcriptional start site (TSS) of the eGene.

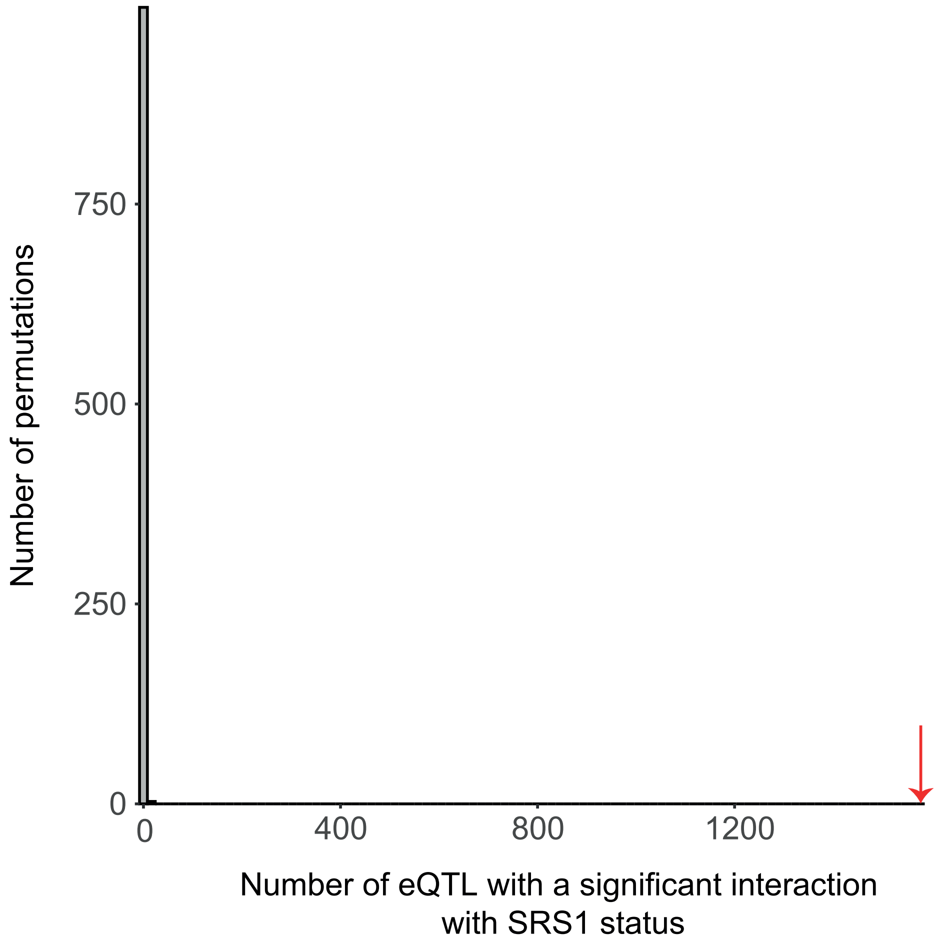


**Figure S10: Distribution of SRS interaction QTL from permutation analysis.**
SRS status was permuted across samples and the eQTL interaction analysis was repeated, with the number of significant interactions for each permutation plotted as a histogram. The observed result is marked with a red arrow.

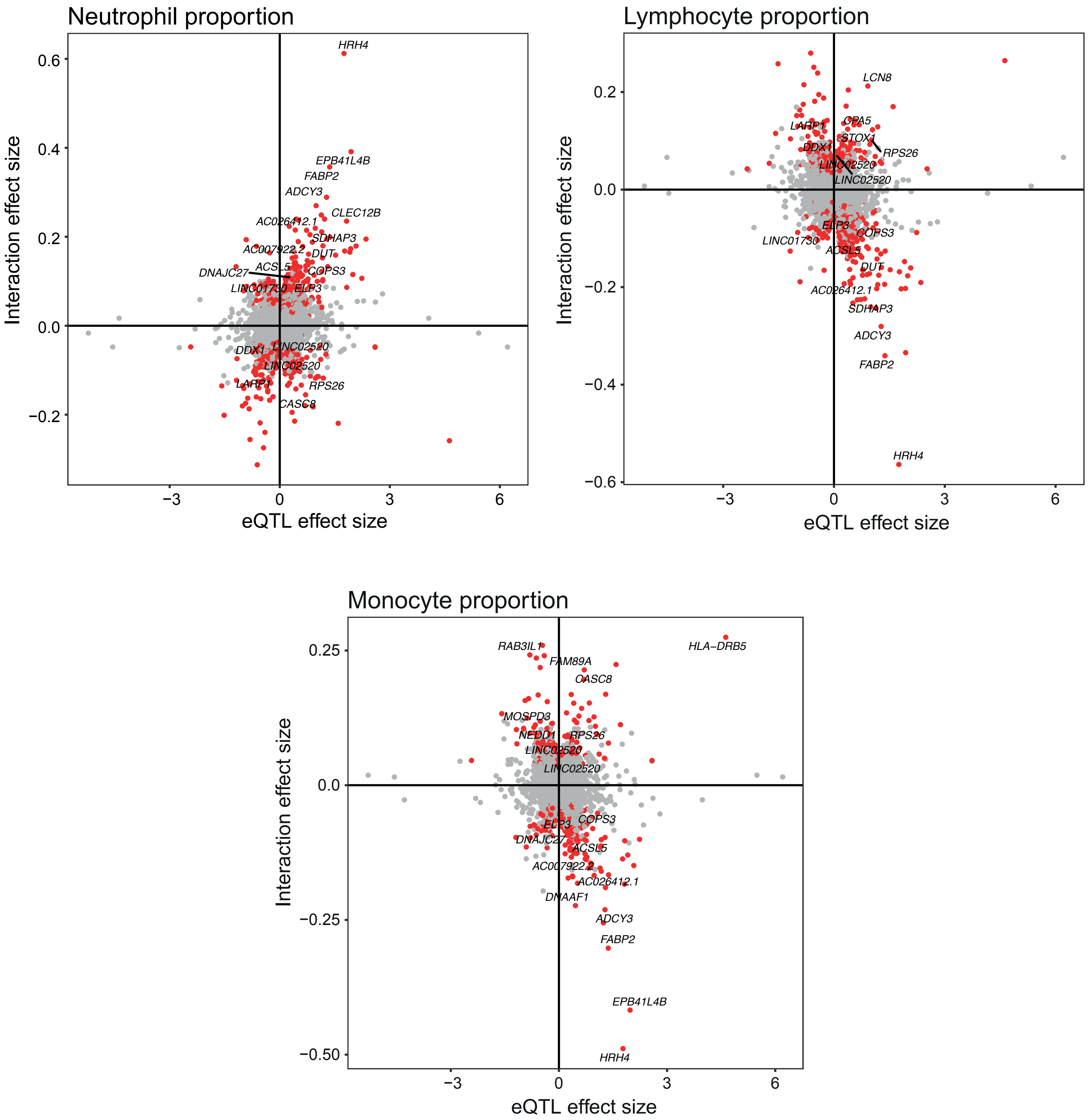


**Figure S11: Cell proportion interaction QTL results.**
eQTL interactions with measured cell proportions. Each point represents an independent eSNP-eGene pair, with the interaction effect size plotted against the genotype effect. eQTLs with bigger effects with increased cell proportion are found in the top right and bottom left quadrants. Red colour indicates a significant interaction between genotype and cell proportion (FDR<0.05), with the most significant results labelled with the eGene name.

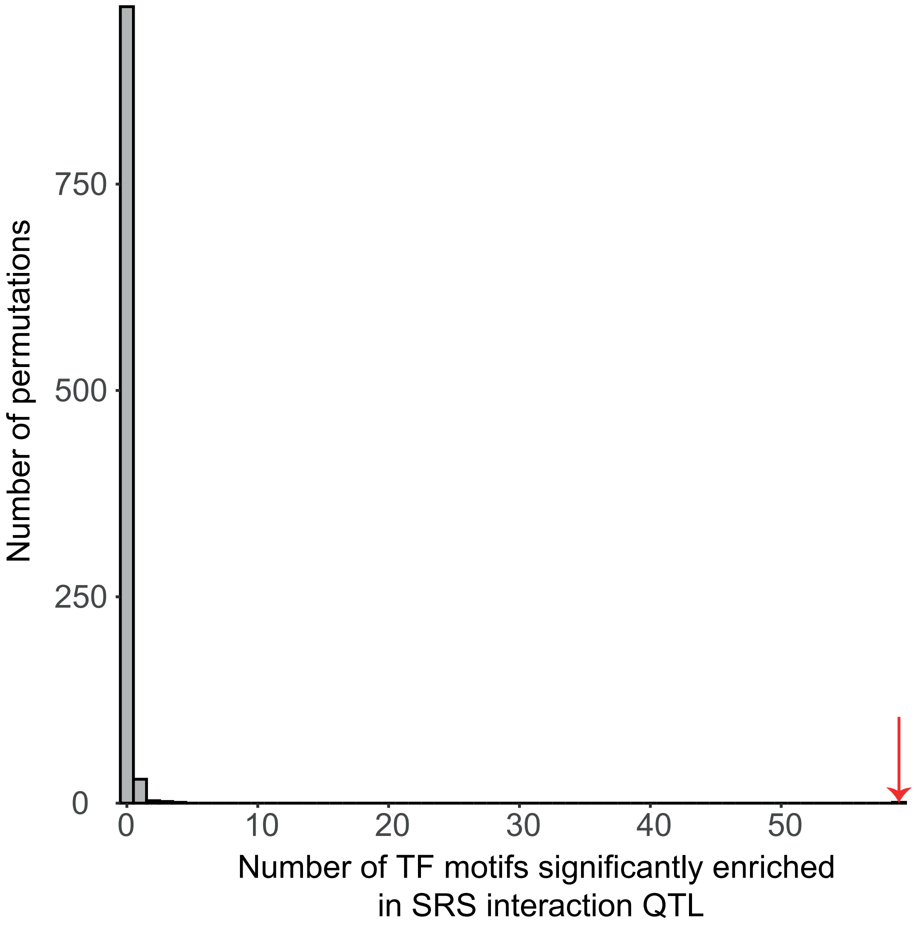


**Figure S12: Distribution of transcription factor binding site enrichment results from permutation analysis.**
eQTL interaction status was permuted across all eQTL signals and the TFBS enrichment tests were repeated, with the number of significantly enriched motifs for each permutation plotted as a histogram. The observed result is marked with a red arrow.

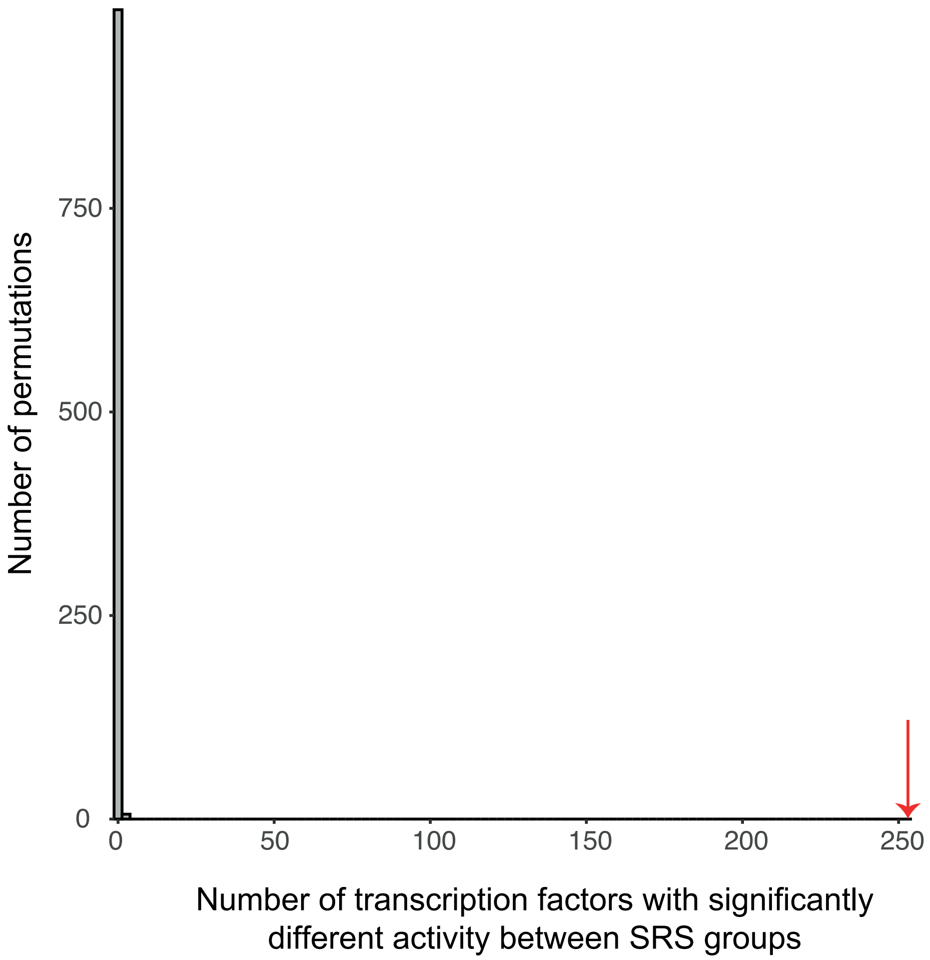


**Figure S13: Distribution of DoRothEA inferred TF activity differences from permutation analysis.**
SRS status was permuted across samples and inferred transcription factor activity compared between groups, with the number of significantly differing TFs for each permutation plotted as a histogram. The observed result is marked with a red arrow.

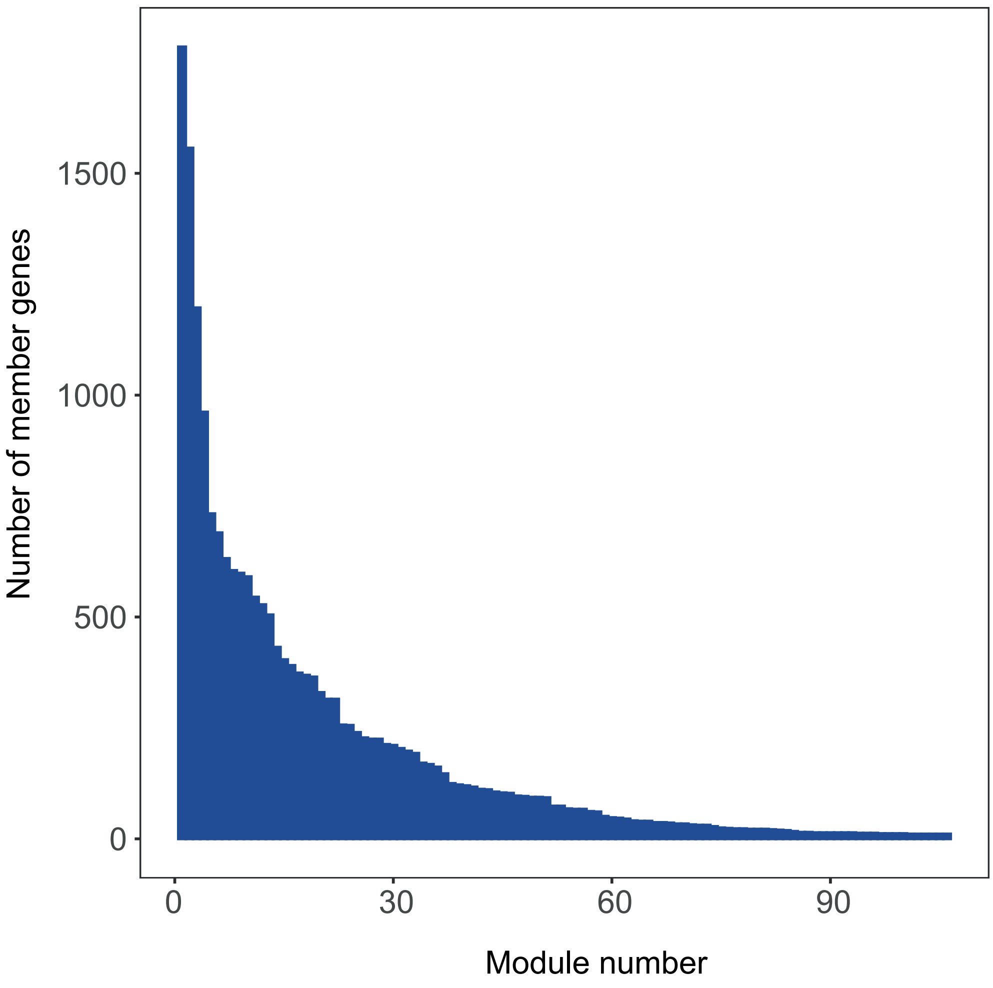


**Figure S14: Histogram of co-expression module size.**
The number of genes in each module ranged from 11 to 1,785.

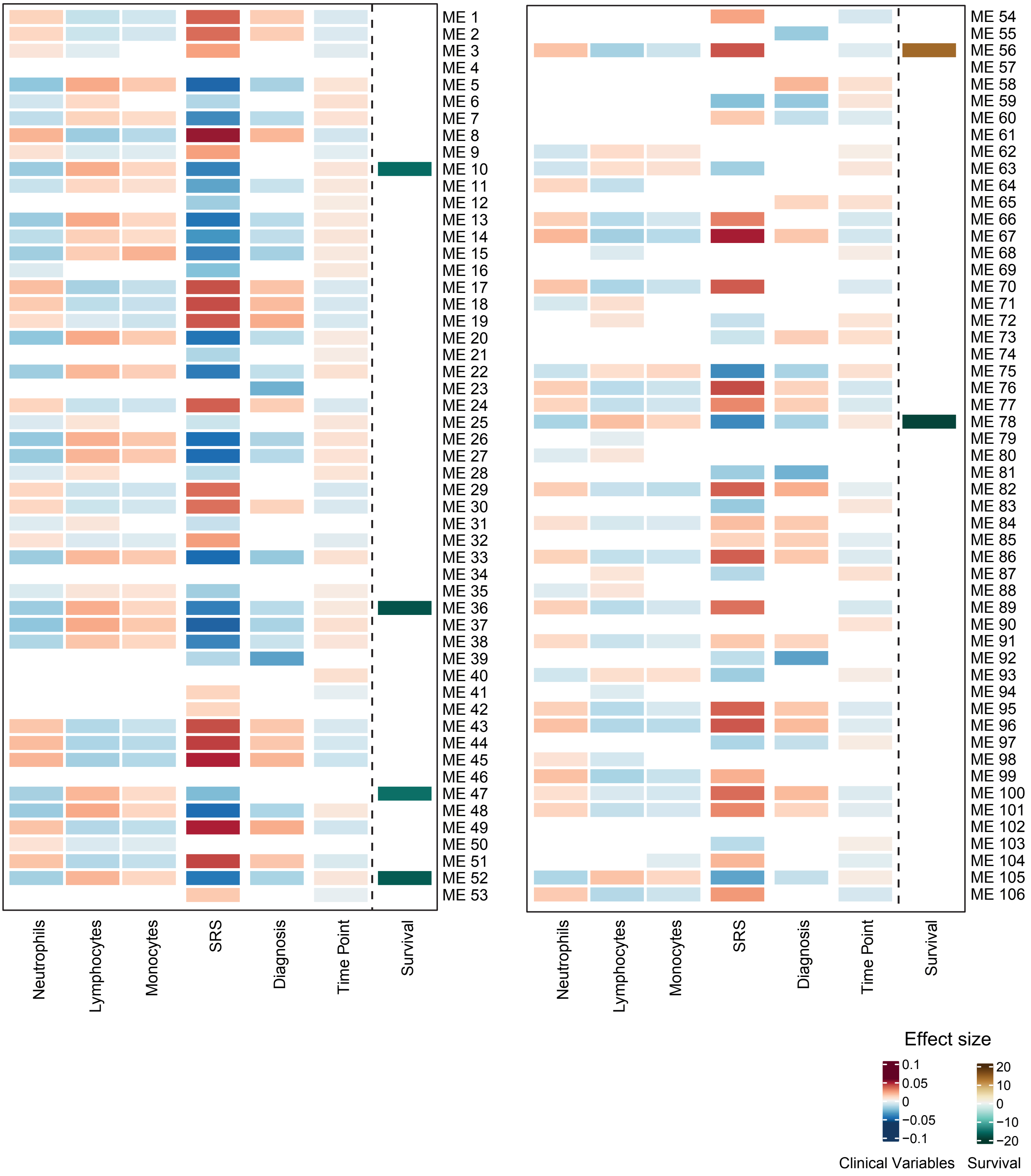


**Figure S15: Module eigengene associations with sepsis endophenotypes.**
Heatmap showing significant associations between all module eigengenes (MEs) and clinical phenotypes. MEs were tested for differential expression with measured cell proportions, SRS1 status, diagnosis (CAP or FP) and time point (day 1, 3, 5) using a linear mixed model. Association of each ME with survival up to 28 days was tested using a Cox proportional hazards model.

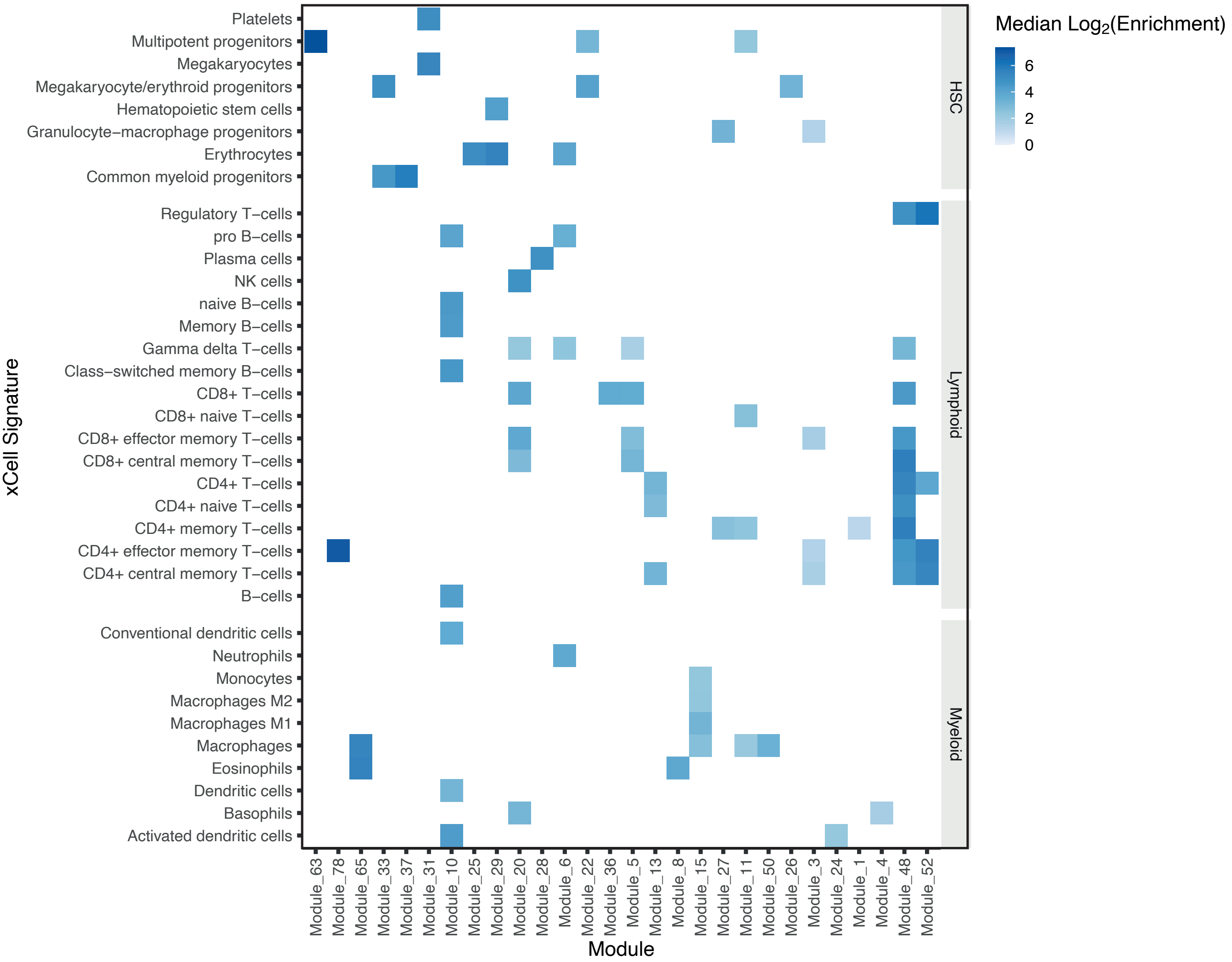


**Figure S16: Enrichment of xCell marker genes in module members.**
Modules were tested for enrichment of xCell gene signatures derived from large whole blood transcriptomic studies. Modules shown had significant enrichment for at least one signature.

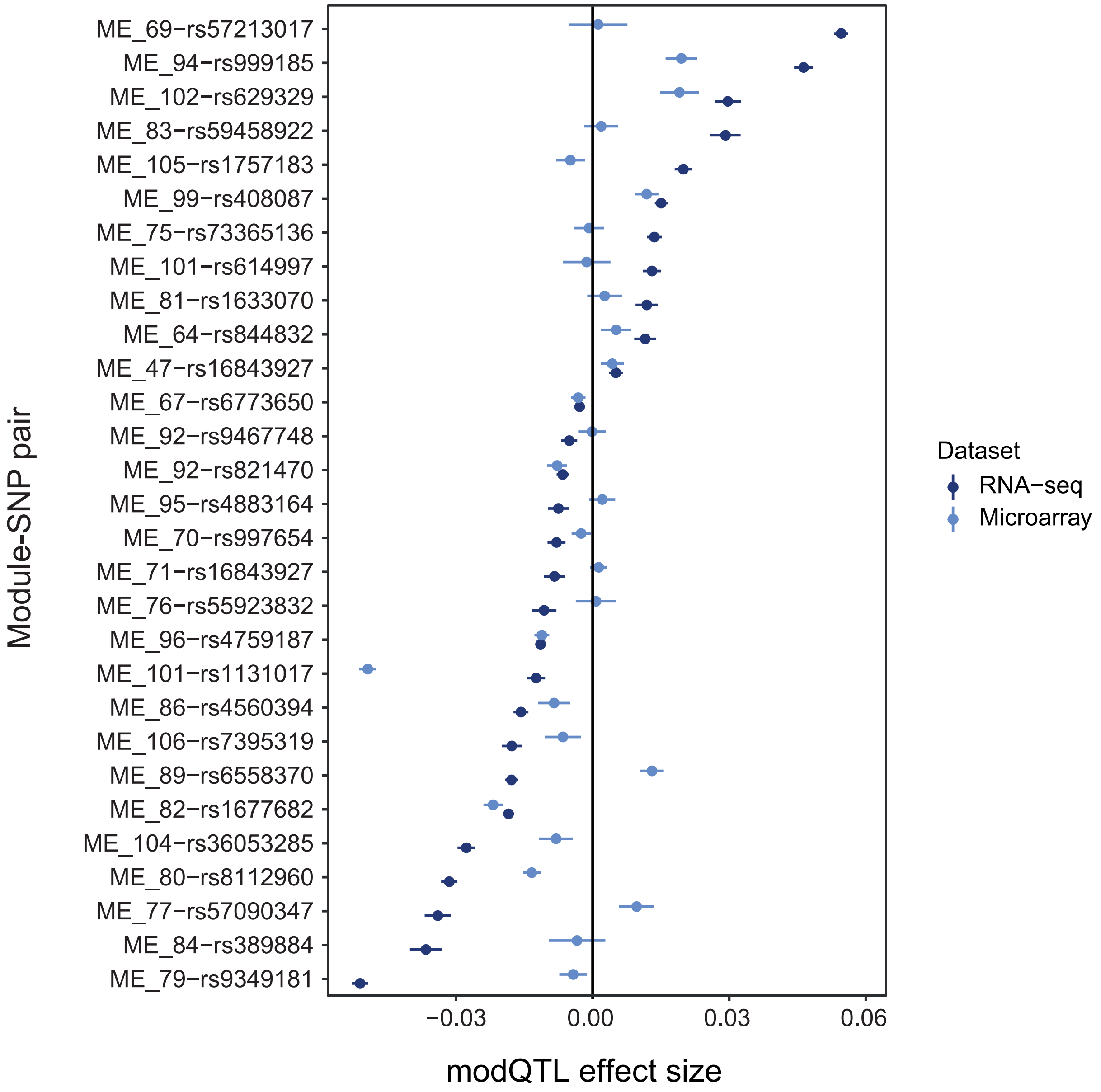


**Figure S17: Replication of modQTL effects in a microarray sepsis cohort.**
Forest plot of modQTL replicated in a validation cohort. Of the 29 modQTL that could be tested, 16 were replicated with consistent direction of effect. Effect sizes from the discovery RNA-seq data set and the replication microarray data set are shown as points with 95% confidence intervals as lines.

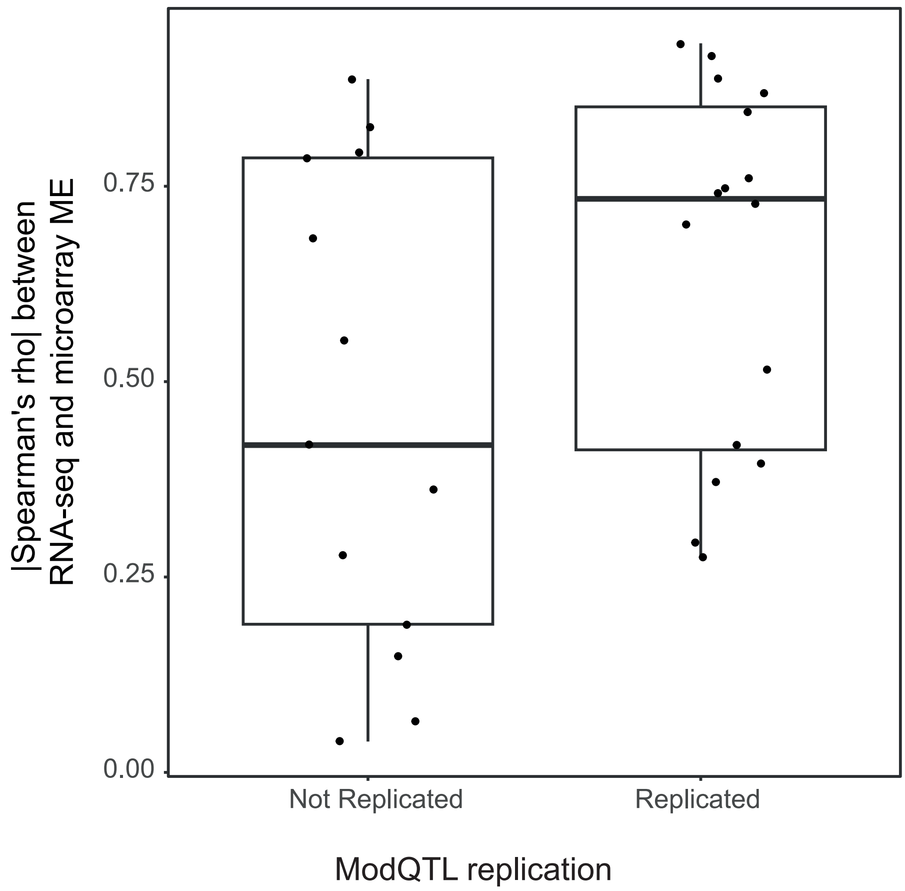


**Figure S18: Correlation of module eigengenes across technologies by replication status.** Module eigengenes were calculated using the same module gene sets in a microarray cohort with 135 overlapping samples. Similarity between module eigengenes were tested using Spearman's rho for the overlapping samples. Module QTL that replicated with the non-overlapping samples had better correlated eigengenes between the two datasets.

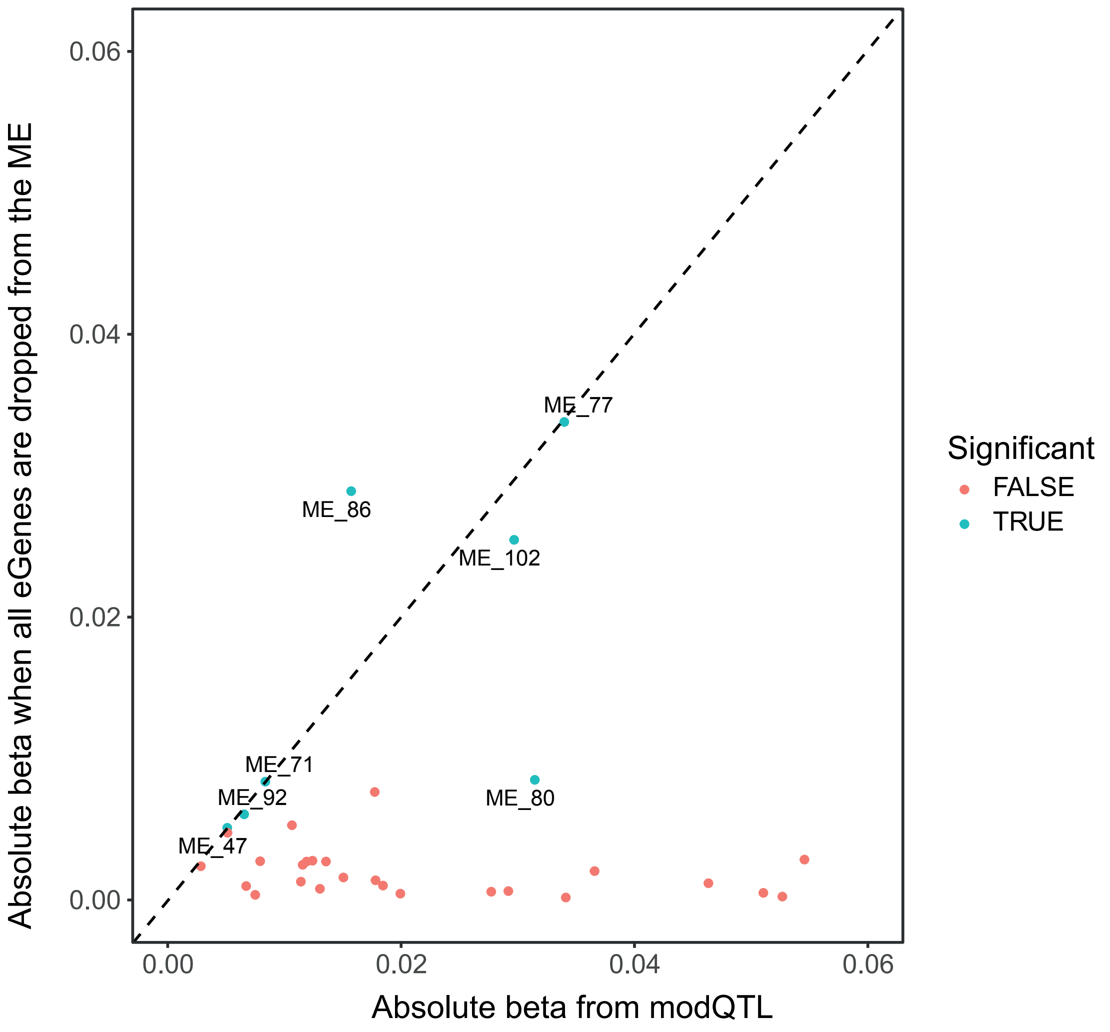


**Figure S19: ModQTL sensitivity analysis.**
Module eigengenes were recalculated excluding any eGenes for the associated modQTL SNPs, and the modQTL retested. The new beta value is plotted against the original for the same SNP-ME pair, and coloured by whether the association remained significant.

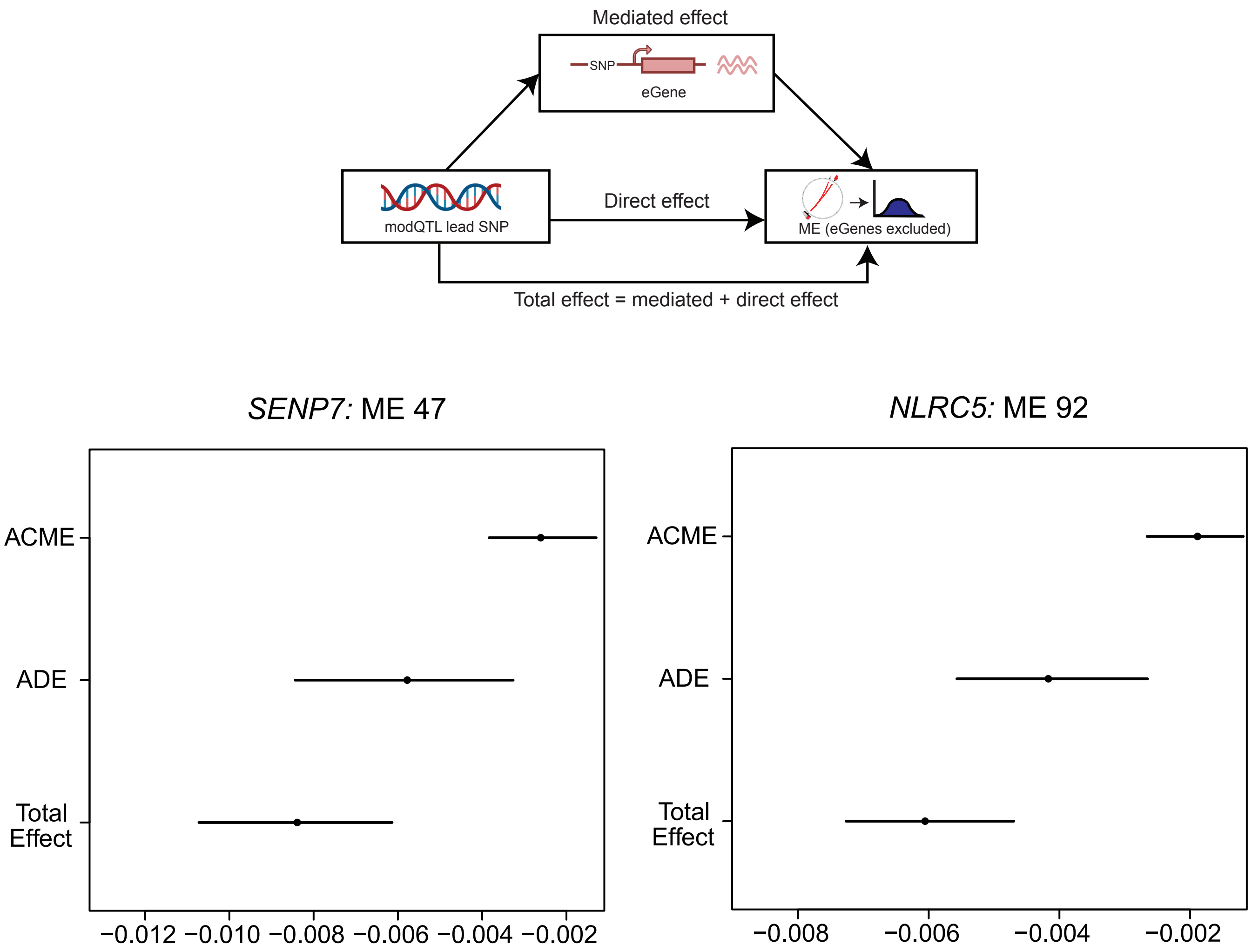


**Figure S20: ModQTL mediation results.**
We tested for mediation of associations between the lead modQTL eSNP and the recalculated eigengene by the modQTL SNPs' target cis-eGene(s). The Average Causal Mediated Effect (ACME), Average Direct Effect (ADE) and total effects are plotted for two modQTL of biological interest.
